# Type 1 diabetes and parasite infection: an exploratory study in the NOD mouse

**DOI:** 10.1101/2024.03.27.586975

**Authors:** Emilie Giraud, Laurence Fiette, Evie Melanitou

## Abstract

Microorganisms have long been suspected to influence the outcome of immune-related syndromes, particularly autoimmune diseases. Type 1 diabetes (T1D) results from the autoimmune destruction of the pancreatic islets’ insulin-producing beta cells, causing high glycemia levels. Genetics is part of its etiology, but environmental factors, particularly infectious microorganisms, also play a role.

It was shown that bacteria, viruses, and parasites, influence the outcome of T1D in mice and humans. We have used the Non-Obese Diabetic (NOD) mouse that spontaneously develops T1D to address the influence of a parasitic infection, leishmaniasis. *Leishmania amazonensis* are intracellular eukaryotic parasites replicating predominantly in macrophages and are responsible for cutaneous leishmaniasis. The implication of Th1 immune responses in T1D and leishmaniasis led us to study this parasite in the NOD mouse model. We have previously constructed osteopontin knockout mice in an NOD genetic background and demonstrated that this protein plays a role in the T1D phenotype. In addition, osteopontin (OPN) has been found i) to play a role in the immune response to various infectious microorganisms and ii) to be implicated in other autoimmune conditions, such as multiple sclerosis in humans and experimental autoimmune encephalomyelitis (EAE) in mice.

We present herein data demonstrating the implication of OPN in the response to *Leishmania* in the NOD mouse and the influence of this parasitic infection on T1D. This exploratory study aims to investigate the environmental infectious component of the autoimmune response, including through Th1 immunity, common to both T1D and leishmaniasis.

## Introduction

Genetic predisposition and environmental triggers are etiological factors for autoimmune conditions. Autoimmune disease incidence is rising in industrialized countries, while in contrast clean environment and vaccination policies control infections. The “hygiene” or “old friends” hypothesis proposes that vaccination and the eradication of infectious microorganisms, in these countries, may contribute to an excessive autoimmune response against self-tissues and molecules by interfering with the immune system’s normal function [1–5]. Supporting this hypothesis, epidemiological data show a North-South gradient of autoimmune disease frequency and opposite geographical distribution to infectious conditions [6, 7].

The evolutionary adaptation between hosts and microorganisms keeps a homeostatic immune equilibrium requiring the presence of both. The Red Queen hypothesis suggests that genetic mutations promote the organisms’ constant adaptation for survival against ever-evolving opposing species. This hypothesis exemplifies antagonistic interactions between parasites and the host allowing for coevolution dynamics to take place [8, 9]. Constant adaptation of the host genome to environmental triggers including infectious microorganisms may be the origin of genomic and/or epigenomic changes required for the host’s survival. Immune function genes are at the forefront for such adaptation and in the absence of infections, the host immune system dynamics may functionally affect immune homeostasis leading to autoimmune, non-self-recognition. Infections play a significant role in the environmental component of the etiology or even protection against autoimmune diseases [10].

Type 1 diabetes (T1D) is an autoimmune condition affecting the insulin-secreting beta cells of the pancreas resulting in hyperglycemia. It represents 7-12% of the total cases of diabetes in the world, estimated at 425 million people worldwide [11]. T1D frequency is higher in urban rather than rural environments [12]. As urbanization expands its incidence increases [11]. Its genetic origin is well established, and over 20 genes are identified as part of the inherited component conferring up to 50-80% of T1D etiology [13, 14]. However, genetics alone cannot account for the rapid expansion of its incidence in the last 100 years [15–18]. Environmental factors including infections by various microorganisms, bacteria, viruses and parasites play a role in modulating autoimmunity. Viral, bacterial, and parasitic infections were reported to elicit or protect disease pathogenesis [18]. A protective role against T1D in the NOD mouse has been reported by the lymphocytic choriomeningitis virus (LCMV) [19, 20], while epidemiological data suggested that enteroviruses such as Coxsackie virus B (CVB) are involved in the initiation or acceleration of the disease in human [20–22]. It was shown that parasitic infections by helminths prevent diabetes and ameliorate insulin secretion [23–25]. Similarly, schistosomiasis protects against diabetes [25] as well as does the TB (Tuberculosis) vaccine against Bacillus Calmette-Guerin (BCG) that induces a host response preventing T1D [26, 27]. The increase of diabetes mellitus in subtropical countries is accompanied by the attenuation of immune defenses and an increase of susceptibility to infections due at least in part to the high levels of most of the time uncontrolled glycemia [28].

Leishmaniasis is a neglected disease group, with a high prevalence worldwide, caused by an intracellular protozoan parasite, *Leishmania spp.* [29]. Diabetes can worsen the outcome of cutaneous lesions caused by *Leishmania (L.) major* or *L. infantum* parasites [30]. Similarly, it was shown that diabetes enhances ulcerated lesions caused by *L. braziliensis* by affecting the pro-inflammatory cytokine levels and impairing response to therapy [31]. *L. amazonensis* (*L. am*.) causes the cutaneous form of the human disease and it is transmitted by the bite of infected phlebotomine sandflies during blood feeding [32–34].

The role of osteopontin (OPN) in the host response to *L. amazonensis* has been evaluated in our previous studies [35]. OPN encoded by the *spp1* gene (secreted phosphoprotein 1) originally identified in activated T cells (also named *eta-1* for early T cell activation-1 gene) is a key cytokine implicated in efficient type 1 immune response (Th1) and differentially regulates *il-12* and *il-10* expression in macrophages (MF) [36]. Knockout mice for the *spp1* gene exhibit severely impaired Th1 immunity to viral and bacterial infections [36]. OPN also is associated with autoimmune diseases, including T1D, in humans and the mouse [37–39]. Moreover, OPN is an early marker for islet autoimmunity in human T1D patients (our unpublished data and [40]). We have previously reported that OPN is involved in the host response to *L. am.* parasites in the C57BL/6 mice [35]. These parasites induce *opn* gene expression in the bone marrow derived macrophages (BMF) *in vitro* and *in vivo* on the sites of parasite inoculation and inhibit the pro-inflammatory transcripts [35].

The role of OPN in the immune system, in both infectious and autoimmune diseases led us to investigate its implication in the infectious environmental component in autoimmune disease etiology, as postulated by the “hygiene” hypothesis. We created mutant mice lacking the osteopontin gene in a NOD genetic background [39] and demonstrated that in its absence T1D is accelerated suggesting a protective effect of OPN on the disease [39].

Our study of the wild-type and knockout NOD mice for the *opn* gene showed that infection with *L. am.* has differential effects on T1D phenotype and parasitic load, depending on OPN expression. We also observed variations in clinical phenotype, parasite content and pro-inflammatory markers in the absence of OPN. After infection with the parasites, a Th1 to Th2 shift was observed in the absence of OPN.

## Results

### OPN-Parasites interactions and T1D

Longitudinal analysis of the cumulative incidence of T1D in the NOD wild (NOD^+/+^) and *opn* knockout mice (NOD.*opn*^-/-^) was carried out in the presence or the absence of *L. am.* parasites (Fig. 1A and B). Survival curves showed that in the absence of OPN (NOD.*opn*^-/-^) T1D is accelerated in comparison with the wild-type animals (Fig. 1A, P<0.0001). In contrast after infection with the parasites, T1D was significantly delayed in the absence of OPN (Fig. 1B, P=0.0260). T1D progression between the infected and non-infected animals was compared taking into consideration the cumulative disease frequencies at two phenotypic windows according to the age, at 15-20 and 21-25 weeks. Although breeding conditions of the NOD colonies influence T1D, usually glycemia appears after 20 weeks of age while most of the animals become diabetic at around 30 weeks [41, 42]. In the absence of OPN, animals develop accelerated diabetes between 15-20 weeks of age (55%), while in contrast at this age, low diabetes incidence was observed in the wild-type animals (20%) (Fig. 1C) [39]. In the presence of the *L. am.* parasites, while the early incidence of T1D (15-20 weeks) remains low in the wild-type animals (33%), the cumulative incidence after 21 weeks increases (83.3% Fig. 1C). In the absence of OPN, *L. am.* infection inhibits T1D of both age groups (33% and 58% respectively) (Fig. 1B and 1C). Moreover, higher mortality at over 26 weeks of age is observed in the infected animals for both wild (60%) and *opn* knockout mice (41%), while in the absence of parasites, the *opn* knockout animals show 0% mortality (Fig. 1D and Table S1A). These data indicate that in the absence of *opn* an immune balance takes place favoring survival, despite high blood sugar levels, whereas in the presence of *Leishmania* a cumulative autoimmune- and infection-related morbidity is taking place (Fig. 1D and Table S1B). However, the waning role of OPN remains unclear. Indeed, while the presence of this protein in non-infected animals, is protective against T1D (Fig. 1C, 40% of NOD^+/+^ versus 70% of NOD.*opn*^-/-^), its absence does not favor an additional deleterious phenotype (morbidity 30% vs 0% respectively) (Fig. 1D). Whilst the evolution of body weight during aging in both WT and KO mice was similar after infection with the parasites (Fig. S1B), lower body weights were systematically observed in the absence of OPN (Fig. S1A). This indicates a contribution of OPN to the general physiological status of the animals while it concurs with the protective role of the OPN against T1D in the NOD mice (Fig. 1A) and as reported previously [39]. However, this effect is abolished in the presence of the parasites as shown by a higher number of diabetic animals (83% in infected vs 40% in non-infected). In the absence of OPN, *L. am.* infection protects against T1D (Fig. 1C and S1C), probably by interfering with the balance of the Th1 responses solicited by the parasite infection. We hypothesized that in the absence of OPN, the Th1/Th2 paradigm of resistance/susceptibility of the host response may shift towards an aggressive for the parasites, Th2 immune phenotype.

**Figure 1:**
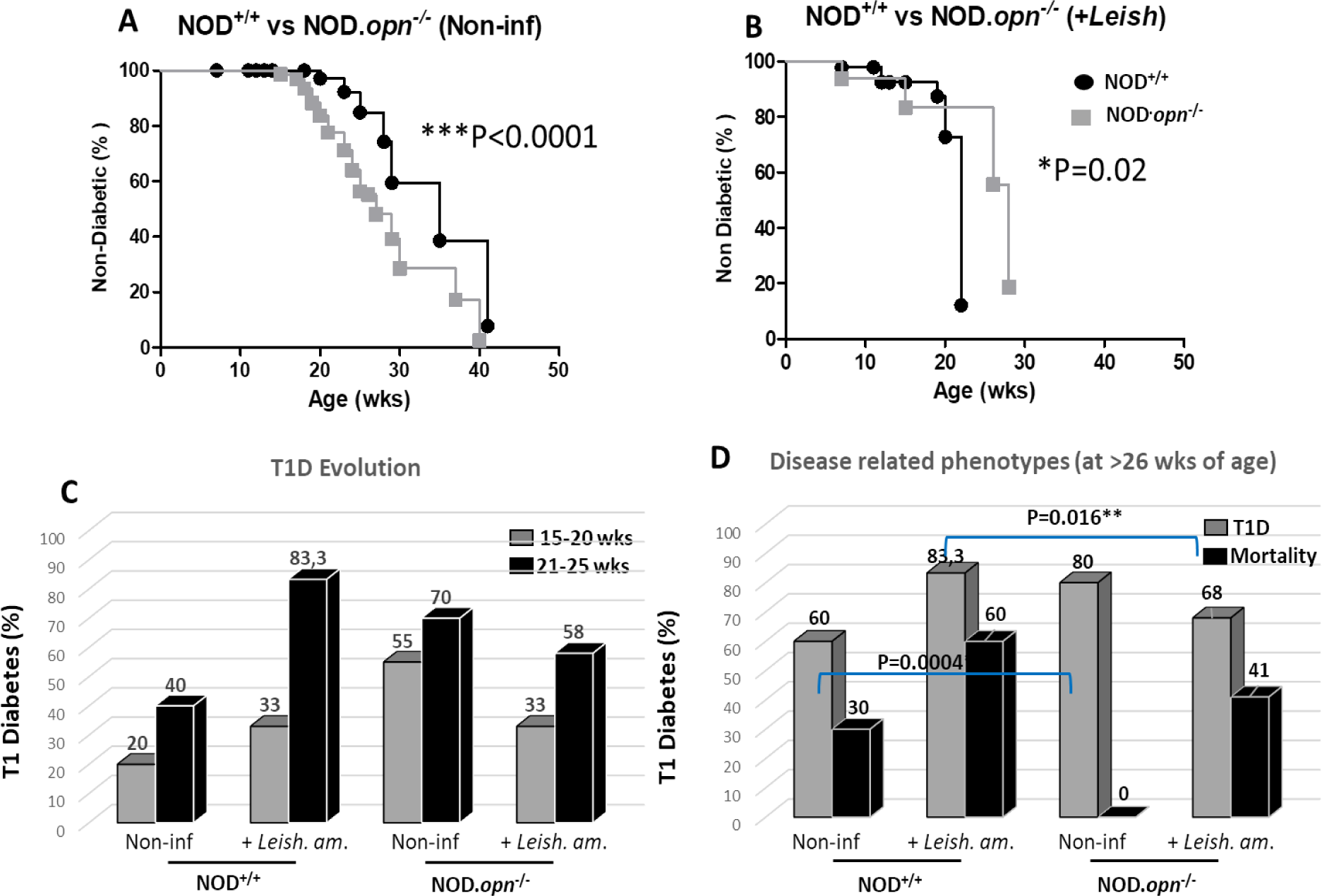
OPN-parasite interactions in T1D phenotype in the NOD mice. **A & B**. Longitudinal analysis of T1D (accumulative incidence, Survival %) in (**A)** non-infected NOD (NOD^+/+^ n°:10) and *opn* mutant (NOD.*opn*^-/-^ n°:20) mice (P<0.0001). (**B)** T1D survival (%) in animals infected with *L. amazonensis* parasites at early pre-inflammatory stages (6-8 weeks). NOD.*opn^-/-^* n°: 12 and NOD^+/+^ INF n°: 6 (P=0,0260). (**C)** T1D evolution (%). Cumulative incidence is calculated at two phenotypic windows: 15-20 weeks and 21-25 weeks of age. (**D)** Disease-related phenotypes, comparative analysis at 26 weeks of age. T1D in NOD^+/+^ vs NOD.*opn*^-/-^ non-infected (P=0.0004) and animals infected with *Leishmania amazonensis* NOD^+/+^ vs NOD.*opn*^-/-^ (P=0.016). (Log-rank (Mantel-Cox) test), (see also Table S1).

### *In vivo* infection of the NOD mice with *L. amazonensis* metacyclic promastigotes

#### 1. Clinical phenotype and parasitic load

To address our hypothesis that the absence of OPN impacts the Th1/Th2 immune balance favoring the persistence of parasites and contributing to lower glycemia, we performed additional *in vivo* experiments but this time by a multiparametric approach to delineate distinct phases after the inoculation of *L. am.*. We assessed the *in vivo* responses against the parasites, of the NOD mice in the presence or absence of OPN. NOD wild-type and *opn* knockout mice were inoculated in the ear dermis with luciferase-expressing *L. am.* metacyclic promastigotes (10^4^), the infectious form of *L. am..* Longitudinal analysis of the clinical phenotype (ear widths) showed variations between the mice with a significant acceleration of the lesion development in the absence of OPN starting at day 32 post-infection (*p.i.)* (Fig. 2A).

**Figure 2:**
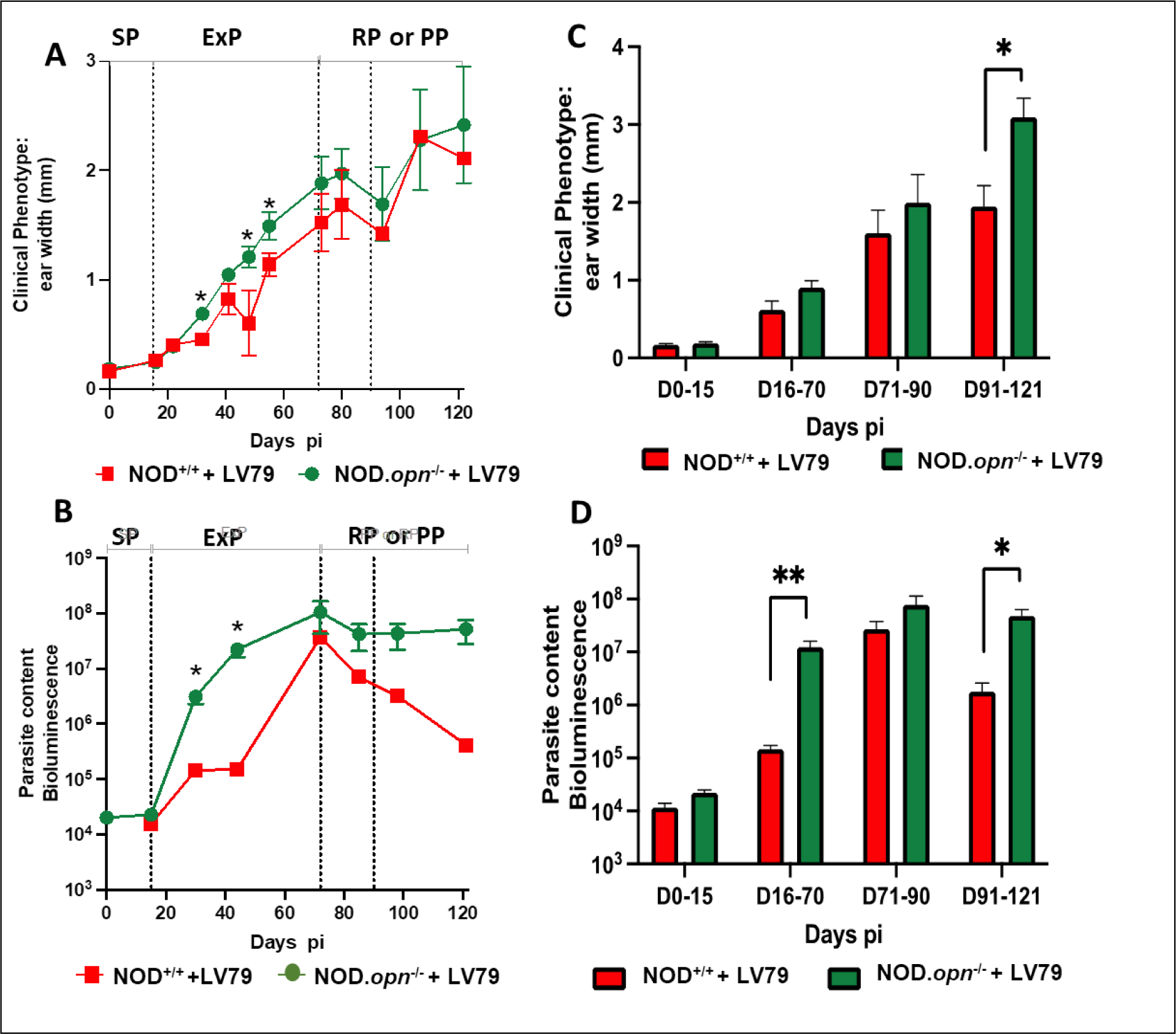
In vivo clinical phenotype evaluation and parasitic load. NOD/LtJ (NOD^+/+^) wild type and *opn* mutant mice (NOD.*opn^-^*^/-^) were inoculated with 10^4^ *L. amazonensis* metacyclic promastigotes in the ear pinna. **(A)** Longitudinal examination of clinical phenotype (ear width) at different time points post-inoculation. Medians and SDs are indicated. Ear widths differences between *opn* knockout and wild-type mice were observed at Day >30 *p.i*. P = 0.0017. The X-axis represents days *p.i.* The four phases of infection (SP, ExP, RP and PP) are described in the text). **(B)** Monitoring of fluctuations of parasite load by bioluminescence. Signals are captured in the dermis of the ear pinnae. Results obtained from wild-type mice (NOD^+/+^: n°: 15 mice per group) are represented in green and from *opn* mutant mice (NOD.*opn*^−/−^, n°: 14 mice per group) are represented in red. **(C)** and **(D)** Time windows (days), after inoculation of 10^4^ metacyclic promastigotes of luciferase-expressing *L. amazonensis* in the ear dermis of wild type and *opn* knockout NOD mice. Animals were grouped for each time window as indicated on the X-axis up to 120 days p.i. (WT: n = 15 mice, KO: n = 14 mice). **(C)** Comparison of clinical lesions at different time points post inoculation (ear widths), (Mann-Whitney test; *P < 0.017). **(D**) Comparison of parasitic load determined by bioluminescence signal quantification at different time points post inoculation (Medians and SD are indicated). (Mann-Whitney test; D16-70 **P =0.0046; D91-121 *P=0.0382).

Consistently, with the clinical phenotype, the parasitic load was higher in the *opn* mutant mice, remaining at high levels after day 70 *p.i.*, while the parasite load of the WT animals significantly decreased (Fig. 2B).

Overall, the clinical phenotype, followed for over 100 days post-inoculation, progressed by four discrete phases (Fig. 2C and D). The first phase (window D0-15 *p.i.*) is a silent phase (SP) whereas neither parasites nor clinical features may be detected (Fig. 2A and B). Then, the expansion phase (ExP) of *L. am.* is associated with an inflammatory aspect of the cutaneous lesion developing into erythematous edema, (window D16-70 *p.i.*). Parasitic load indeed increased over time in both wild-type and *opn* mutant mice and this increase was significantly higher and appeared earlier in the absence of OPN (Fig 2B and D). Then the parasitic load reached a peak (window D71-90 *p.i.*). The last phase is either a reduction phase (RP) for the WT mice, associated with a significant reduction of parasite load (Fig. 2B and D), or a plateau phase (PP) for the KO mice, associated with the maintenance of parasite load at a high level (window D91-121 *p.i.*). Histological evaluation of the infected ear tissues showed discrete differences in the microscopic lesions between the wild type and the *opn* mutant mice at 100 days *p.i.* corresponding to the last phase (Fig. 3A and 3B). Parasite content was higher in the absence of osteopontin as well as macrophage numbers (Fig. 3D and Fig. S2C & D). At this time point, despite the rather discrete differences in the impact of the *Leishmania* parasites in both wild-type and *opn* mutant mice, the histological examination of the infected sites confirmed the higher content of parasites (Fig. S2D) as well as the pronounced ulceration in the *opn* knockout mice (Fig. 3C), while necrosis was more pronounced in the presence of *opn* (Fig. 3C). Differences of the impact of parasite infection related to OPN were evident at the earlier phases after infection (Fig. 2A and 2B).

**Figure 3:**
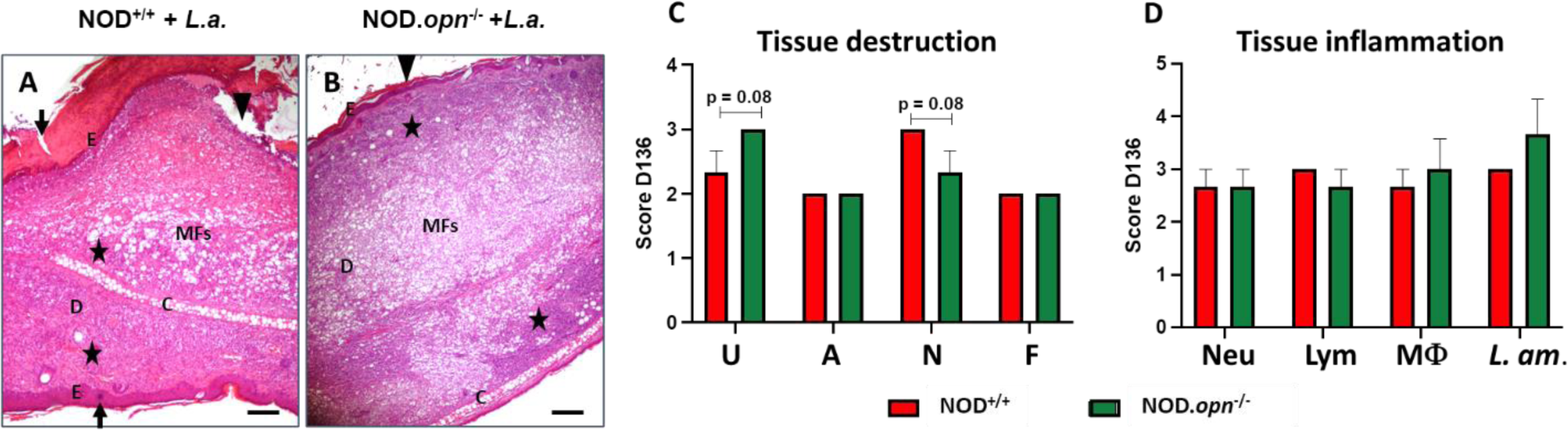
Histopathology and evaluation of the tissue inflammation markers on the site of parasite inoculation. **(A)** Histological features of ear lesions in wild (NOD^+/+^, n°: 5) and **(B)** *opn* mutant mice (NOD.*opn*^-/-^, n°=6) at 136 days *p.i*. (age 25 weeks). Microscopic lesions showed discrete differences between wild-type and knockout mice: parasites and MF content were higher in the absence of osteopontin. Triangles: ulceration, *stars: inflammation, infected macrophages, arrow: acanthosis: E: epidermis, D: dermis, C: cartilage. Hematoxylin and Eosin (H&E), original magnification x4, scale bar: 100 µm. **(C) and (D).** Histological scores from NOD^+/+^ (red bars) and NOD.*opn*^-/-^ mice (green bars), post-inoculated with 10^4^ *L. amazonensis* metacyclic promastigotes. **(C).** Tissue destruction and **(D).** Tissue inflammation. **(C) and (D)** aabbreviations: U= ulceration, A= acanthosis, N= necrosis, F= fibrosis, Neu= neutrophils, Lym=lymphocytes, MF=macrophages, *L. am*.= *L. amazonensis.* Statistics 2way Anova.

#### 2. In vivo host response to the parasites and role of OPN: Comparative analysis between the NOD and C57BL/6 mice

In the C57BL/6 genetic background, we have previously shown an increase of the inflammatory lesions after *L. am.* infection in the ear pinna of the *opn* knockout in comparison with the wild mice, while in contrast, the parasite content remained similar [35].

As mentioned above, in the NOD mice, four phases were delineated: a silent phase (SP), an expansion phase (ExP), a pick phase (PP) and a reduction (RP) (in the WT mice) or a plateau phase (in the KO mice) (Fig. 2). Taking into consideration the Th1-related autoimmune particularities of the NOD mice [43], and the absence of *Leishmania* data for this strain, we compared the *in vivo* parasite responses between the two strains in the presence and the absence of OPN.

Parasite load and ear lesions showed few differences between these two strains (Fig. 4A and B). In wild mice parasite content and ear width were lower in the NOD than in the C57BL/6, especially between 16-70 days post-infection (expansion phase) (Fig. 4A and B), while at the plateau phase (D91-121) the ear lesion was more pronounced in the NOD mice (Fig. 4B).

**Figure 4:**
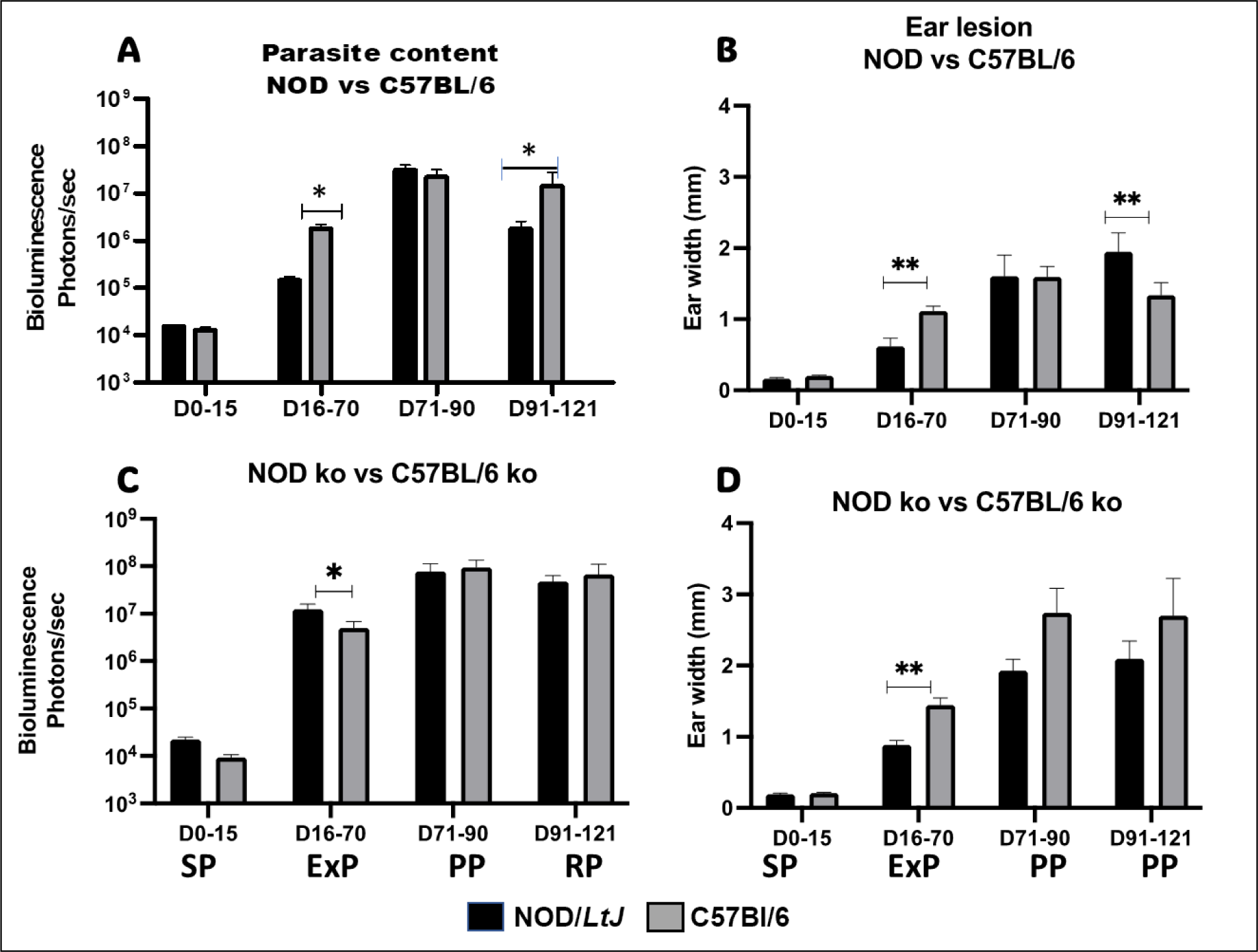
Comparative analysis of *Leishmania amazonensis* infection between NOD and C57BL/6 mice in the presence or the absence of osteopontin. Cumulated data of four phenotypic windows are represented corresponding to days post-infection. **(A) and (C).** Matched evaluation of the fluctuations of parasitic load by bioluminescence signaling (expressed in photons/sec/ear), in sites of inoculation *in vivo* of NOD/*LtJ* (black bars) and C57BL/6 (grey bars) strains. **(A):** wild type, D16-70 (*P=0.0469); D91-121 (*P=0.0382) and (C): *opn* knockout mice, D0-15, (***P=0,0001) and D16-70 (*P=0.0196). **(B) and (D)** Ear lesions (ear widths expressed in mm) **B:** of wild-type NOD and C57BL/6 mice D16-70 (*P=0,0196); D91-121 (**P=0.0028) and (**D)** of *opn* knockout animals; D16-70, (**P=0.0005). A Mann-Whitney test was performed to compare the fold changes between the two groups. Non-parametric correlation tests (Spearman) were performed to compare Bioluminescence.

In contrast, in the *opn* knockout mice, the parasitic load was consistently higher and similar between the two strains except for the ExP phase whereas NOD^-/-^ showed higher parasitic load (Fig. 4C). Additionally the ear lesions of the NOD KO mice were less developed than the C57BL/6 KO mice (Fig. 4D). These data indicate that the Th1 related immune nature of the NOD genome facilitates the proliferation of the parasites in the absence of *opn*, at least at the early phases (Fig.4C), while at its presence (Fig. 4A) these responses are moderated probably by the presence of synergistic effects exerted on the *opn* by the infection and the autoimmune-prone genetic setting in relation with the Th1/Th2 balance.

Host cellular responses to *L. amazonensis*: *in vitro* infection of BMF isolated from wild and *opn*-mutant NOD mice.

*L. am.* amastigotes, are predominantly found within the resident dermal macrophages [44] and in dendritic cells [45] in the mammalian host. We evaluated the response to infection by *L. am.* parasites and the implication of OPN, in BMF isolated from the NOD mice.

We first evaluated the replication of parasites at the amastigote stage (the intracellular stage) in BMF, at 24h and 48h *p.i.* in the presence and the absence of OPN (Fig. 5).

**Figure 5:**
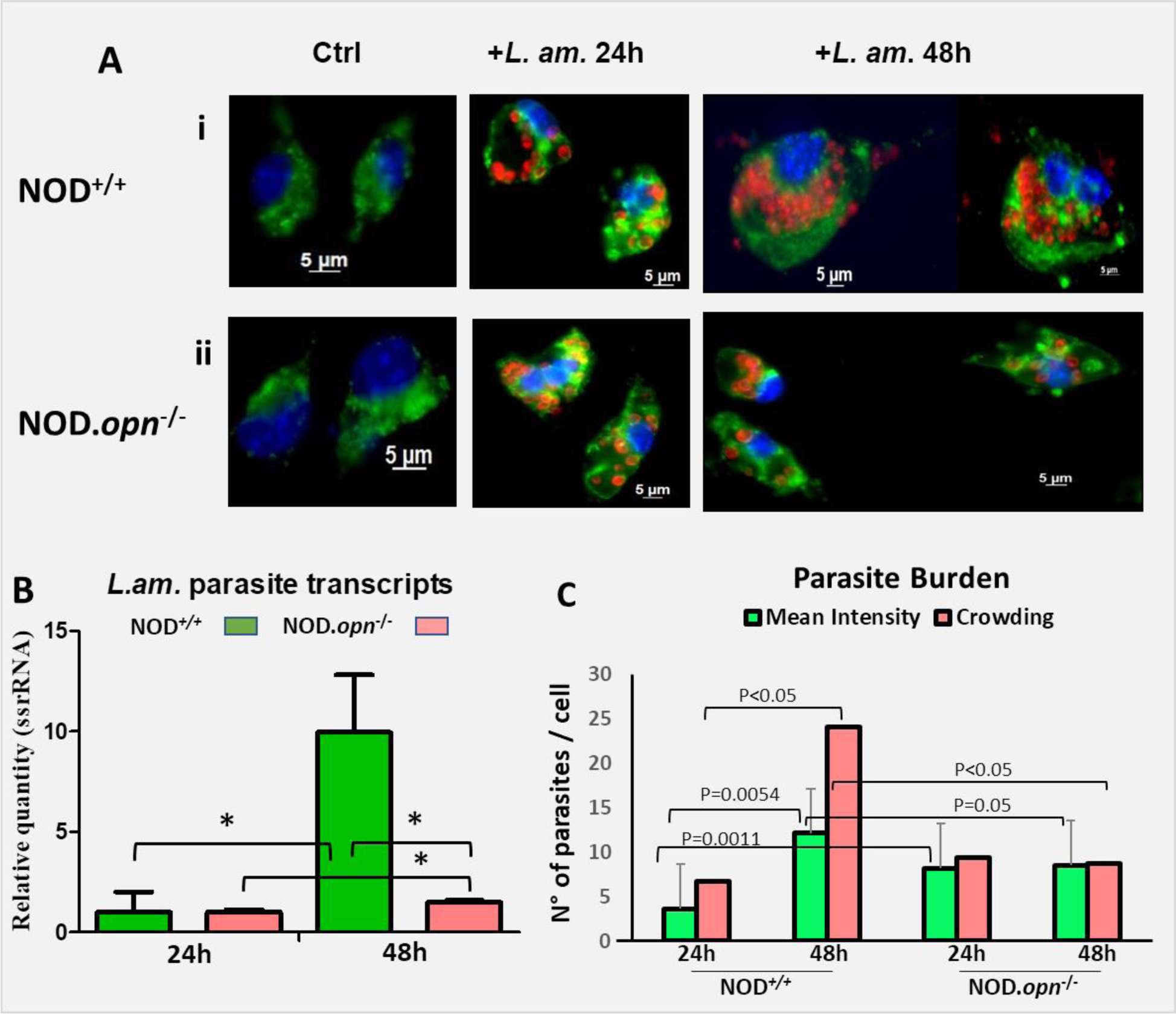
*L. amazonensis* proliferation in BMFs in the presence or absence of osteopontin in the NOD genetic background. **(A)** BMFs isolated from NOD/*LtJ* wild-type **(i)** or *opn* mutant (NOD.*opn*^-/-^ mice (**ii**) were infected with *L. amazonensis* amastigotes at a ratio of 4:1. Representative images of BMFs populations are shown as well as control non-infected BMFs (ctrl). Twenty-four and 48 hours later, each BMF population was analyzed by immunostaining. Nuclei were stained with Hoechst (blue), vacuoles with Lysosome-associated membrane protein Lamp-1 Ab and FITC-labelled conjugate (green), and amastigotes with 2A3-A26 Ab and Texas Red-conjugate (red). All images are taken in phase-contrast optical microscopy. **(B)** Parasite quantification by Q-RT-PCR of the relative expression of ssrRNA of *Leishmania amazonensis* in the cells in the presence or the absence of osteopontin, as evaluated at 24 h and 48 h post-infection. Statistics: Unpaired t-test with Welch’s correction: WT 24h vs WT 48h: P=0.0364; WT 48h vs KO 48h: P=0,0363; KO 24h vs KO 48h: P value= 0,0155. **C.** Parasite Burden in BMF. Mean intensities and intracellular parasite crowding of *L. amazonensis* amastigotes per infected cell were monitored by manual analysis of immunofluorescence image captures (AxioVision) of at least 2 different experiments (average number of 60 evaluated cells per condition). Statistical analyses were performed using the QP3.0 program designed for Quantitative parasitology as described in the Methods section. Mean intensities were compared by the Bootstrap test and 2-sided bootstrap p-values are given as follows: WT vs KO at 24 h, P=0.0011; WT 24 h vs 48 h P=0.0010 and KO 24 h vs 48 h, NS. Mean crowding was significant for WT 24h vs 48h P<0.05, Cl 97.5%, and WT 48h vs KO 48h P<0.05 Cl 97.5%. Statistics are analytically presented in Tables S2A and B)

Immunostaining of BMF isolated from the NOD wild mice infected with the parasites showed increased proliferation of the *Leishmania* amastigotes at 48h when compared to 24h *p.i.* (Fig. 5A). *Opn* KO macrophages were similarly infected by the parasites (Fig.5A), however no or minor proliferation was observed between 24h and 48h *p.i.* in the absence of OPN (Fig. 5A). qRT-PCR of the *ssrRNA* of *Leishmania* confirmed these data (Fig. 5B). Thus, in the NOD genetic background osteopontin seems to participate in parasitic proliferation (Fig. 5B and Table S2).

Parasite burden evaluation, (mean intensity and crowding) of *Leishmania*, followed the same patterns observed by immunostaining and qRT-PCR and confirmed these data (Fig. 5C and Table S2). Mean intensities and parasite crowding were significantly higher (P<0.05) in the wild mice at 48h *p.i.* (12.13 and 24.07 parasites/cell respectively) than in the KO mice (8.5 and 8.76 parasites/cell respectively) (Fig. 5C and Table S2A). Contrary to the NOD, in the BMF isolated from C57BL/6 *opn* knockout mice, higher infectious rates were observed in comparison with the wild-type cells indicating that OPN is involved in cell protective responses against the parasites in this strain [35]. However, while 100% of the NOD KO macrophages were infected with the parasites at 24h, the wild-type macrophages reached 90% cell infection only at 48h *p.i.* (Fig. S3).

These data are consistent with the implication of OPN in the adaptation of *Leishmania* parasites in their cellular niche, the BMF of the NOD mice, possibly participating in the Th1 parasite-elicited immune responses.

### *L. amazonensis* parasites stimulate *opn* gene expression and OPN protein in BMF isolated from NOD mice

The ability of parasites to multiply in the NOD BMF, in the presence of OPN rather than in its absence, indicates that this protein is required at least for their proliferation in the host macrophages. The effect of infection by *Leishmania* amastigotes on the osteopontin gene expression was evaluated by immunostaining and qRT-PCR (Fig. 6A and 6B respectively). A four-fold increase of *opn* transcripts in the presence of parasites, relative to non-infected macrophages, was observed at 48h *p.i.* (Fig. 6B). The density of the OPN protein was evaluated in the infected cells and compared with the non-infected control cells (Fig. 6C). OPN protein follows similar patterns at 48h *p.i.*, however at 24h *p.i.* an increase of the protein also is observed (Fig. 6C).

**Figure 6:**
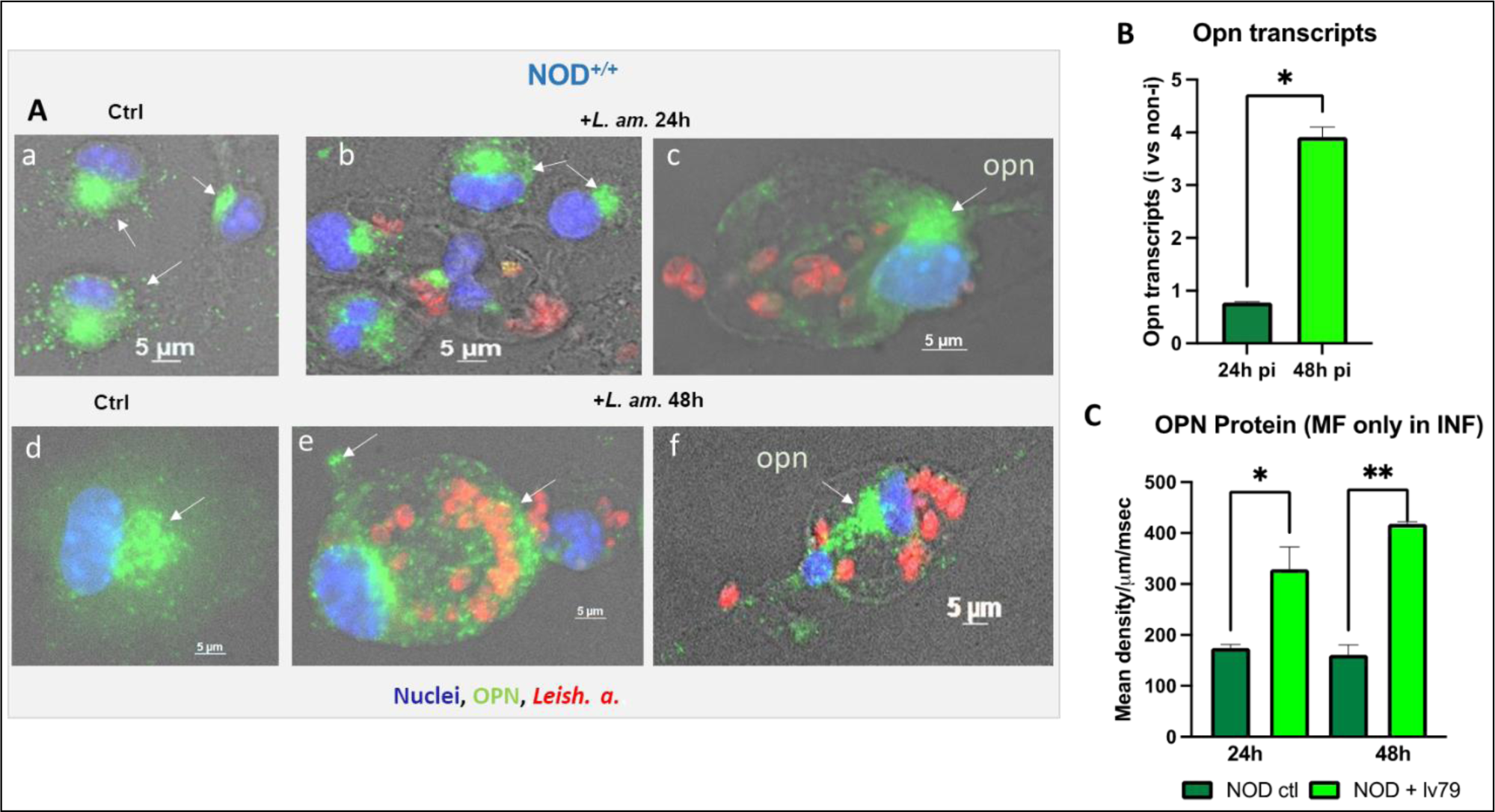
OPN expression in BMF of the NOD^+/+^ mice, in the presence of *Leishmania amazonensis*. **(A)** Immunofluorescence staining of BMFs. **a** & **d:** control non-infected at 24h and 48h respectively. **b** & **c**: BMFs infected with LV79 at 24h and **e** & **f** at 48h post-infection. Green: OPN, red: LV79; blue: nuclei. **(B)** qRT-PCR of OPN transcripts (*P=0.0294). **(C)** OPN protein quantification from control non-infected (ctl) and infected with +LV79 BMFs. NOD ctl 24h vs NOD+lv79 24h (*P=0.0454); NOD ctl 48h vs NOD+lv79 48h (**P=0.0091). Mann Whitney test, two-tailed.

These data together with the lack of parasite proliferation in the NOD.*opn*^-/-^ mice, described above (Fig. 5B and 6B), confirm that the endogenous presence of osteopontin in the NOD BMFs is implicated in the host response to parasite infection. The Th1 immune environment of the NOD mice and the Th1 cytokine properties of the OPN seem to favor the *Leishmania* parasites within the NOD BMF niche, whilst, in contrast, the presence of OPN confers protection against early T1D onset only in the absence of infection (Fig. 1A).

This seems to be OPN-dependent since the evaluation of CD44 transcripts, encoding for the OPN receptor remained unchanged in both wild-type and *opn* mutant mice (Fig. S4) suggesting a receptor-independent effect of the OPN molecule. Therefore, OPN favors parasite proliferation in the BMF *in vitro* (Fig. 5), while *in vivo* the presence of OPN seems to be host-protective against the *Leishmania* parasites as seen in the clinical phenotype (Fig. 2C) and the parasite content in the infected tissue (Fig. 2C and D). Different mechanisms and/or molecules implicated *in vitro* in the BMF and *in vivo* in the infected tissue may explain these observations.

The *L. am.* effect on the *opn* transcripts (Fig. 6B) was compared between the NOD, DBA/2 and BALB/c mice (Fig. S5). It is to be noted that the BALB/c mice are sensitive to *Leishmania* parasites while the DBA/2 are resistant [46]. Interestingly, *opn* transcripts were induced by the presence of the parasites in the BALB/c macrophages, while NOD mice showed lower *opn* gene expression and DBA/2 intermediate values at 24h *p.i.* Therefore, *Leishmania*, in a permissive sensitive genetic background (BALB/c mice) stimulates early *opn* gene expression (24h *p.i*.), that in its turn facilitates parasitic proliferation in the BMFs (Fig. S5). Overall, the variation of the *opn* gene expression in response to *L. am.* observed between these three strains outlines i) the implication of OPN in parasite proliferation independently to the genetic susceptibility to leishmaniasis disease and ii) a strain-specific response to the parasites requiring the OPN, together with possible additional molecules participating to the variation of the host response to the parasites.

### OPN favors parasite proliferation in macrophages in the NOD genetic background

To further evaluate the response to parasite infection of the NOD mice, we assessed the role of OPN in cell infectivity and survival by immunostaining BMF cells infected with *Leishmania* parasites at 24h and 48h *p.i.* in the presence or the absence of OPN (Fig. 7 and Fig. S6). OPN is expressed in both intracellular (iOPN) and extracellular (sOPN) forms in the BMF (Fig. 7A, i & ii). High parasite proliferation was observed in the macrophages at 48h *p.i.* in the presence of OPN (Fig. 7B vii and Fig. S6, ii, iv, v). The cells show intact, non-apoptotic nuclei (Fig. 7 and Fig. S6 i and ii). In contrast, the number of parasites is lower in the NOD *opn* knockout mice (Fig. S6 iii and vii). We have identified a similar phenotype in *Leishmania-*infected BMF isolated from C57BL/6 mice, but only in the absence of OPN [35]. Following these data, the inflammatory response to the parasites is contained as shown by the modulation of *il-1β* transcripts at 48h *p.i.* in the presence of OPN as well as in its absence (Table 1.1). However, the two-fold increase of *il-1β* transcripts at 24h *p.i*. in the NOD wild mice indicates an initial host response to the parasites, also dependent on the presence of OPN as confirmed by the downregulation of *il-1β* transcripts at its absence (Table 1.1). Similar data were obtained in the ear pinna, after *L. am.* infection *in vivo* (Fig 8). Il-1β is a key mediator of the inflammatory response to pathogens [47]. *Leishmania spp* parasites effectively down-regulate *il-1β* in the C57BL/6 mice [35]. The increase of *il-1β* transcripts at Day 100 *p.i.* may indicate the presence of a local reservoir of parasites after the infection is contained, as previously reported for *L. major* [48]. Together, these observations suggest i) that in the macrophages isolated from the NOD mice, OPN favors the growth of the amastigotes and ii) the parasites while eliciting the host response as shown by the increase of *il-1β* transcripts at 24h *p.i.*, confer an adaptation to the host, as seen by the subsequent downregulation of *il-1β* at 48h *p.i*. Overall, the NOD genetic background seems to represent a parasite-favorable cellular environmental niche.

**Figure 7.**
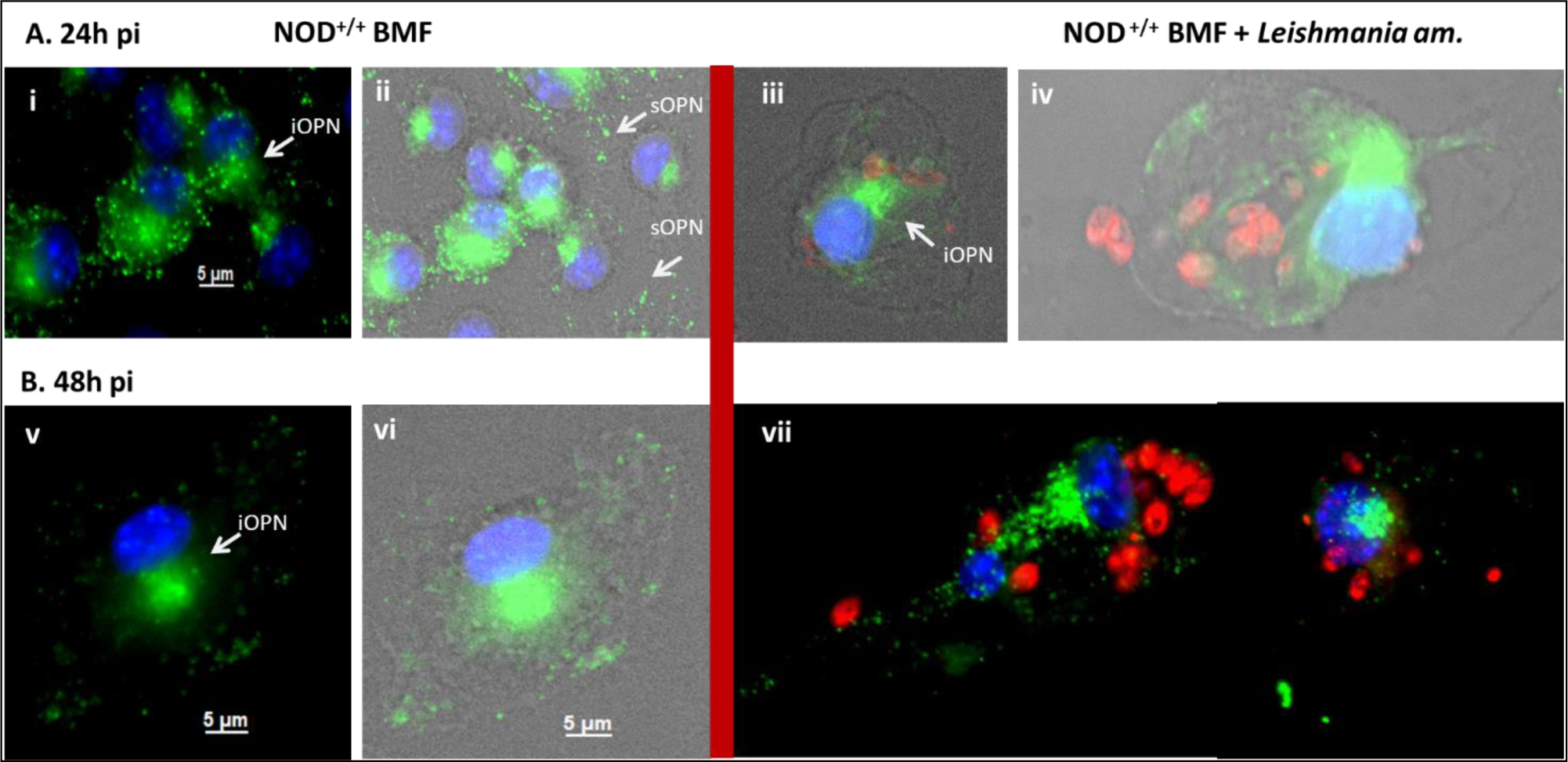
Infectivity and survival of *Leishmania amazonensis* parasites in the BMFs isolated from the NOD mice. Representative images of immunofluorescence staining for OPN protein (green) and *Leishmania* parasites (red), nuclei are in blue. **(A)** BMFs at 24h in culture non-infected (i and ii) and infected (iii and iv). **(B)** BMFs at 48h in culture non-infected (v and vi) and infected (vii). Total N° of cells at 24h: WT: 117; KO: 29 and at 48h *p.i.* WT: 55; KO: 4. iOPN: intracellular OPN; sOPN: secreted OPN. The image was captured by Zeiss AxioVision Rel. 4.8.2 image acquisition software (Carl Zeiss International).

**Figure 8.**
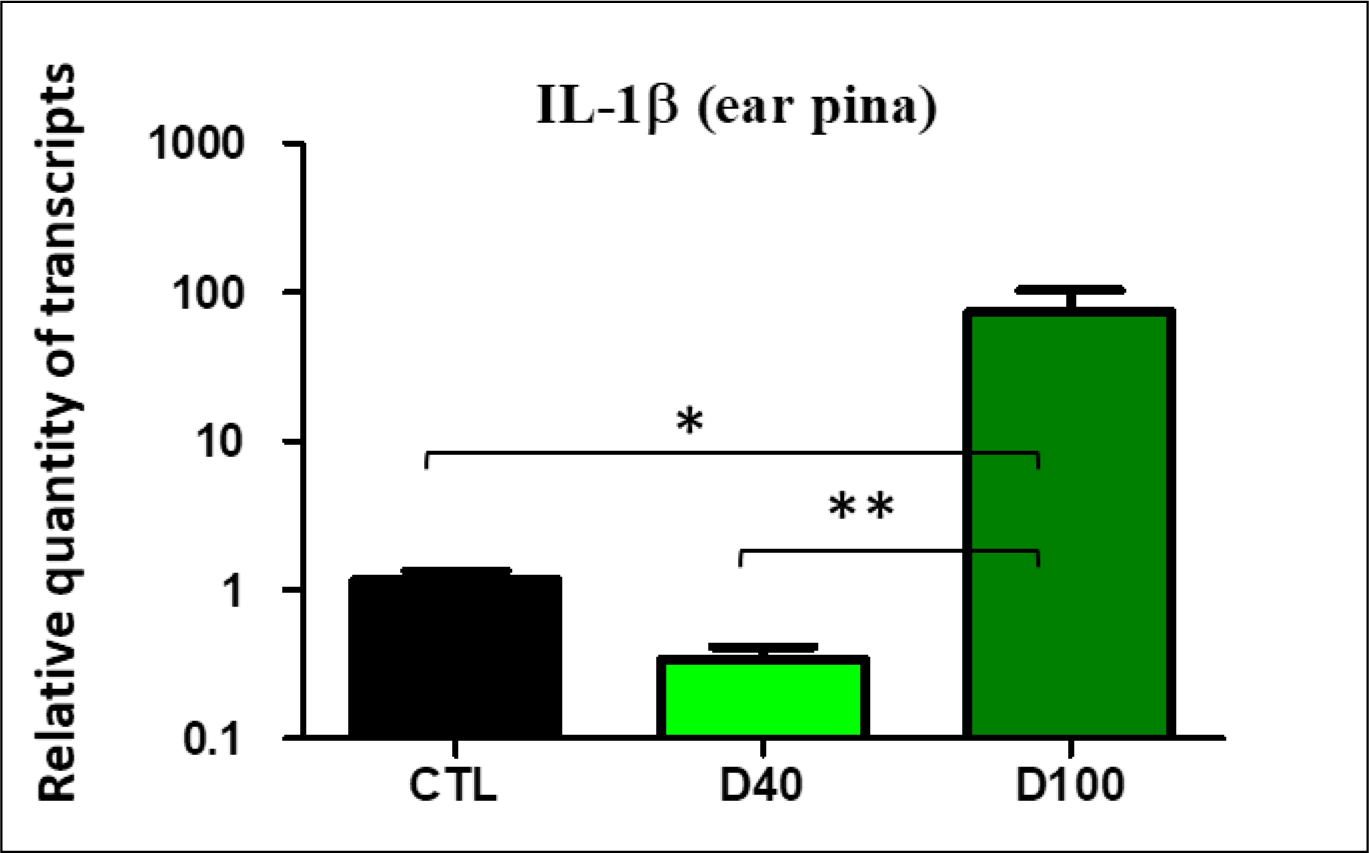
***In vivo*** q**RT-PCR** from total RNA isolated from the infected ear pinna tissue of WT (CTL) infected, D_40_ *p.i.* and KO mice at D_40_ and D_100_ *p.i.*. (Ctl *vs* D_100_ *p.i.*: P=0,0051** and KO D_40_ *vs* KO D_100_ *p.i.*: P<0.0001). F-test to compare variances.

**Table 1.** Host response to *Leishmania amazonensis* and OPN role in immune responses. 1.1. *In vitro* Transcript profiles encoding for il-1β and for iNOS, stat1, il-17 (Th1) and il-10, il-4 (Th2) cytokines in BMF isolated from NOD^+/+^ (WT) and NOD.*opn*^-/-^ (KO) mice in response to *L. amazonensis*. Cells were inoculated with *Leishmania amazonensis* amastigotes at a MOI: 4:1 (parasites: macrophages) and incubated for 24h and 48h before lysis for RNA preparation. Mean fold changes for transcripts and standard deviations have been calculated by using as a calibrator the wild-type non-infected value at each time point.

| Transcripts | BMF | 24h *p.i.* | 48h *p.i.* |
| --- | --- | --- | --- |
| *il-1β* | WT | 2.17 ± 0.2 | -1.42 ± 0.18 |
| | KO | -1.2 ± 0.06 | -1.47 ± 0.05 |
| *iNOS* | WT | 64.44 ± 2.475 | 185.3 ± 3.182 |
| | KO | 32.50 ± 1.768 | 30.50 ± 1.768 |
| *stat1* | WT | 6.077 ± 0.01453 | 14.09 ± 0.2991 |
| | KO | 2.977 ± 0.5488 | 1.593 ± 0.1386 |
| *il-17* | WT | -15.22 ± 0.5691 | -149.5 ± 0.9780 |
| | KO | 18.46 ± 0.1125 | 20.97 ± 0.5000 |
| *il-4* | WT | -1.9 | -2.6 |
|  | KO | -2.55 | -2.62 |
| *il-10* | WT | 0.7700 ± 0.02 | 12.35 ± 0.44 |
| | KO | 21.15 ± 0.16 0.7700 | 56.16 ± 0.17 |

### Immune Host Response to *L. amazonensis* and the role of OPN

L. amazonensis stimulates iNOS expression by a mechanism OPN-dependent.

It was demonstrated that microbial compounds and LPS activate NO in macrophages through the enzymatic action of iNOS (inducible nitric oxide synthase) which represents one of the major microbicidal mechanisms of macrophages against pathogens [49–51]. The antimicrobial role of NO in the control of infectious parasites has been demonstrated in macrophages [52–55], including in the control of *Leishmania* proliferation [56]. Regulation of the inhibition of *Leishmania* parasite growth is mediated by feedback of NO production by iNOS through the action of OPN.

Osteopontin, induced by NO down-regulates iNOS by an autocrine mechanism upon stimulation of the ubiquitin (Ub)-proteasome degradation factor which inhibits stat1 (signal transducer and activator of transcription 1), the critical transcription factor for iNOS gene expression [57]. However, *Leishmania* parasites can also act directly on stat1. It was demonstrated that *L. major* and *L. mexicana* attenuate stat1 and *L. donovani* infection of macrophages attenuates tyrosine phosphorylation of JAK1, JAK2 and stat1 molecules, subsequently blocking IFN-γ and thus evading the host defense mechanisms [58, 59].

We have addressed the effect of the parasites on the iNOS transcripts in the BMF in the presence or absence of OPN (Table 1.1). In the BMF of the NOD wild mice, iNOS transcripts are highly upregulated at 24h *p.i.,* reaching 185-fold after 48h *p.i.* (Table 1.1), while in non-infected cells, iNOS remains low (data not shown). This up-regulation is inhibited in the absence of OPN (Table 1.1).

We addressed the transcript levels of *stat1*, known to stimulate iNOS gene transcription. While *stat1* transcripts showed 6 and 14-fold increase (at 24h and 48h *p.i.* respectively) in the wild-type BMF, low quantities were observed in the KO cells indicating the implication of OPN (Table 1.1). Stat1 is known to act as a dimer after phosphorylation and nuclear translocation, to trigger the transcription of its targets [60]. The activation of *stat1* in the BMF (14-fold at 48h, Table 1.1) correlates with the increase of iNOS. However, parasite burden is higher in the NOD wild mice despite the high iNOS gene expression (Fig. 5B), while in the absence of OPN parasite burden is lower despite the decrease of the iNOS transcripts, indicating the necessity of the presence of OPN for parasite proliferation (Fig. 5C). The presence of iNOS together with the sustained presence of the parasites in the NOD BMF is unexpected unless another mechanism is involved containing the host response to the parasites and related with the presence of OPN.

Therefore, despite the known iNOS effect on pathogen control, it seems that *Leishmania* parasites evolved additional complex adaptation mechanisms to dampen the host defense responses. Two hypotheses may be proposed for these observations: i) in the NOD mice, NO is not acting directly to contain the parasites; other mechanisms seem to be implicated and ii) insufficient presence of oxygen may dampen NO production, despite high iNOS, resulting in no clearance of the parasites. A similar phenomenon was demonstrated in macrophages infected with *L. major* whereas the absence of oxygen failed to produce enough NO to clear *L. major* [61].

The downregulation of iNOS *in vivo*, at the expansion phase (ExP) of the parasite (D_40_), in the sites of inoculation (Table 1.2) and the draining lymph nodes (DLN) (Table 1.2) of the *opn* knockout mice, as well as the increase of iNOS, observed at D_100_ *p.i.* (reduction (RP) phase) correlates with the low *stat1* expression (Table 1.2) and may indicate parasite persistence in tissue reservoirs as previously demonstrated [62]. Additional studies are required to confirm and elucidate these observations.

**1.2.** In vivo. Transcript profiles as indicated in the ear pinna and draining lymph nodes (DLN) of NOD^+/+^ (WT) and NOD.*opn*^-/-^ (KO) mice in response to *L. amazonensis*. 10^4^ metacyclic promastigotes of parasites (LV79 strain) were inoculated in the ear dermis of the mice and transcript profiles were evaluated in the temporal windows spanning the early and expansion phase (EP-ExP) at Day_40_ *p.i.* and the parasite reduction phase (RP) at Day_100_ *p.i.* (see Fig. 2). Mean fold changes for transcripts and standard deviations have been calculated in the *opn* KO mice by using as a calibrator the wild-type D_40_ *p.i.* value. Significant differences (p<0.05*) are indicated (Mann-Whitney test or Kruskal Wallis test).

| Transcripts | Tissue | ExP-D<sub>40</sub> (KO/WT)<br>M ± SD | RP-D<sub>100</sub> (KO/WT)<br>M ± SD |
| --- | --- | --- | --- |
| *iNOS* | Ear | 0.42 ± 0.073 * | 46.83 ± 33.28 * |
| | DLN | 0.9167 ± 0.2437 | 13.50 ± 12.20 |
| *stat1* | Ear | 0.79 ± 0.1302 | 2.14 ± 0.95 |
| | DLN | 0.72 ± 0.38 | 0.41 ± 0.06 |
| *il-17* | Ear | 0.013 ± 0.0033 | 104.5 ± 100.2 |
| | DLN | 85.86 ± 78.72 | 1481.34 ± 979.8 |
| *il-4* | Ear | 0.33 ± 0.058 | 15.18 ± 14.59 |
| | DLN | 3.45 ± 3.5 | 30.79 ± 25.1 |
| *il-10* | Ear | 2.533 ± 0.9543 | 578.1 ± 547.8 |
| | DLN | 1.003 ± 0.44 | 0.66 ± 0.05 |

## 2. Role of OPN on the Th1-Th2 immune balance of the host response to *L. amazonensis*

Osteopontin, as mentioned above, is a Th1 cytokine expressed in cells of the adaptive (T cells) and innate (macrophages, DCs) immune system [63]. We evaluated by qRT-PCR the transcripts encoding for genes prompting Th1 or Th2 host immune responses to *L. am.* parasites in the NOD mice in the absence or in the presence of osteopontin *in vitro,* in the BMF (Table 1.1). *Leishmania* infection highly inhibited interleukin-17 (*il-17*) transcripts in the NOD BMF (Table 1.1). Similar data were obtained for the *il-4* transcripts (Table 1.1). However, in the absence of osteopontin, *il-17* was about 20-fold up-regulated (Table 1.1) while *il-4* remained low (Table 1.1). In *Leishmania* susceptible BALB/c mice, *il-4* was reported to be elevated after infection and this leads to progressive disease [64, 65]. The downregulation of *il-4* in the NOD mice indicates that this strain is resistant to the parasites.

In contrast, *il-10* transcripts were induced by the parasites in the presence of osteopontin but only after 48h *p.i.* (12-fold) (Table 1.1). Interestingly, *il-10* transcripts were over 50-fold induced at 48h *p.i.* by the parasites in the BMF of the *opn* knockout mice (Table 1.1). Similar data were obtained *in vivo* in the infected ear pinna of the NOD *opn* mutant mice but only at 100 days *p.i.* (Table 1.2). These data indicate that in the NOD mice, the Th1 inner immune proinflammatory environment is contained by the parasites while in the absence of OPN, the increase of *il-10* transcripts (Table 1.2) indicates a counterbalance towards a protective for the parasites’ host environment and parasite proliferation. The increase of *il-10*, a Th2-related cytokine, in the macrophages (Table 1.1) correlates with the inhibition of parasite proliferation in the presence of OPN *in vivo* (Fig. 2B and D). Il-17 was shown to promote the progression of cutaneous leishmaniasis (CL) in the BALB/c mice known to develop Th2 immunity and succumb to infection [66]. In contrast in the NOD mice, known to possess Th1 immunity, *il-17* is inhibited by the parasites in the BMF (Table 1.1). Increased levels of *il-17*, as well as *il-10* in the absence of OPN *in vitro* and *in vivo*, seem to be under the control of the infection by the parasites and correlate with the post-infection acceleration of T1D (Fig. 2B). Indeed, it was described that NOD mice deficient for il-10, were protected from organ-specific autoimmunity [67], therefore in our experimental setting, the increase of *il-17* and *il-10* transcripts in the *opn*-deficient mice correlates with the T1D acceleration observed after infection by the parasites. Overall, these data are concordant with the increased proliferation of the parasites in the wild-type BMF (Fig. 5) indicating host adaptation of the parasites in the presence of OPN.

## 3. Low expression of IFN-γ in the absence of OPN infers to a Th2 shift promoted by *L. amazonensis* infection

The presence of IFN-γ in Th1 immune cells and the role of this cytokine in preventing the shift to Th2 immunity [68] [69] prompted us to assess its presence in the ear lesions of the infected NOD mice. In the presence of the parasites, IFN-γ transcripts were low in the ear lesions in both wild-type and KO mice (Table 1.3), while in the DLN, a fivefold increase was observed in the presence of opn (Table 1.3) indicating IFN-γ producing cells were present in the DLN after infection with the parasites. The ratio of *IFN-γ* : *il-10* remained low in the absence of OPN on the sites of parasite inoculation (Table 1.3), while the *IFN-γ* : *il-4* ratio showed a 3-fold increase (Table 1.3). However, at Day_100_ *p.i*., both ratios (*IFN-γ*: *il-10* and *IFN-γ* : *il-4*) remained low indicating a shift towards a Th2 response in the absence of OPN and correlating with the high levels of *il-4* and *il-10* observed in the ear lesions as well as in the DLN (Table 1.2). These data concord with published observations showing that *IFN-γ* critically regulates *opn* gene expression. It was reported that in human monocytes while OPN positively regulates *IFN-γ* expression, this cytokine in turn stimulates OPN expression in a positive regulatory loop, during infection [70]. Therefore, in our setting, in the absence of OPN, *IFN-γ* expression is low, and the parasites create a permissive environment, Th2-dependent, especially at D_100_ *p.i.* whereas an increase of Th2 transcripts profiles, is observed (*il-4* and *il-10*, on Table 1.2).

**1.3.**
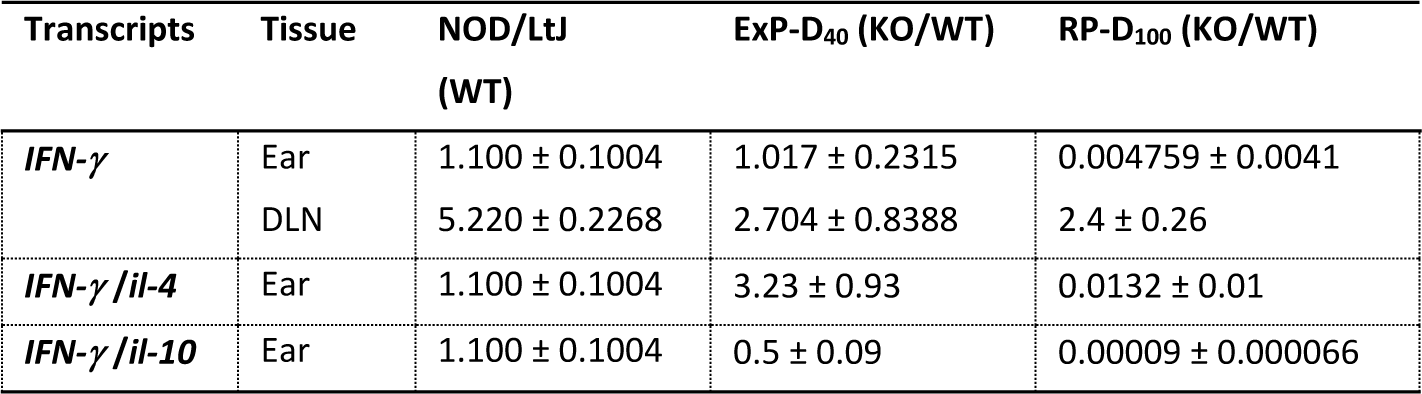
Th2 parasite-dependent shift delineated by low *IFN-γ* expression in the absence of OPN.

| Transcripts | Tissue | NOD/LtJ<br>(WT) | ExP-D<sub>40</sub> (KO/WT) | RP-D<sub>100</sub> (KO/WT) |
| --- | --- | --- | --- | --- |
| *IFN-γ* | Ear | 1.100 ± 0.1004 | 1.017 ± 0.2315 | 0.004759 ± 0.0041 |
|  | DLN | 5.220 ± 0.2268 | 2.704 ± 0.8388 | 2.4 ± 0.26 |
| *IFN-γ/il-4* | Ear | 1.100 ± 0.1004 | 3.23 ± 0.93 | 0.0132 ± 0.01 |
| *IFN-γ/il-10* | Ear | 1.100 ± 0.1004 | 0.5 ± 0.09 | 0.00009 ± 0.000066 |

The role of IFN-γ in *Leishmania* infections has been long debated. In the absence of IFN-γ, C57BL/6 mice are highly susceptible to *L. major* with an expansion of Th2-type responses [71], while two months after infection with *L. am.* they develop late severe disease [71]. Similarly, the implication of IFN-γ in T1D is not clear, ranging from no effect to disease exacerbation [71, 72]. In recent work, loss of IFN-γ-signaling affects T1D incidence but does not prevent it [73]. The acceleration of T1D in the absence of OPN after infection with the parasites and the proliferation of *L. am.* observed in the knockout mice is concordant with a shift towards a Th2 phenotype resulting in exacerbation of the clinical *Leishmania* phenotype (Fig. 3). These observations are in agreement with the low levels of *IFN-γ* observed in the absence of OPN while the parasites delay T1D appearance (Fig. 1) and aggravate the inflammatory clinical phenotype (Fig. 2).

In the absence of OPN the increase of the *IFN-γ* : *il-4* ratio in the *opn* knockout animals *in vivo* in the ear lesions at D_40_ *p.i.* (Table 1.3), indicates that a Th1/Th2 balance is taking place, probably related to a wound repair mechanism which is dependent upon the presence of macrophages and other lymphocytes producing IFN-γ [74]. This suggests rudimentary conservation of Th1 immunity after infection with *Leishmania* parasites, even in the absence of OPN. Yet further analysis is necessary to confirm these data and the proposed hypothesis.

## Discussion

Evolutionary constraints imposed by infectious microorganisms on their hosts impact the adaptation of both the microorganisms and their mammalian hosts. These environmentally imposed cohabitations led to mutualism, commensalism, or parasitism. In the case of parasites, the remnants of such adaptation are present in both the mammalian hosts and the parasites. The adoption of the macrophages as a niche for *Leishmania* parasites is one remarkable example whereas both the host cell and the parasite have found mechanisms for coexistence.

Genetic changes have been fixed throughout evolution conferring adaptation possibilities between parasites and their hosts. The “Hygiene” hypothesis includes the possibility that such genetic changes may be etiological triggers of, or protective against autoimmunity. For example, the control of CL as revealed by genetic studies, depicted a complex picture dependent upon the genetic background of the animals under study. Genetic loci influence the host’s response to the parasites [75, 76]. H2 loci were found to influence the outcome of the infection and depending on the HLA alleles, mice are classified as resistant or prone to CL [77–79]. Similarly, HLA loci and in particular *HLA*-DR/DQ haplotypes are associated with autoimmune diabetes [80]. According to the H2 haplotypes, NOD mice predisposed to T1D carrying the H2^g7^ K^d^ alleles may be classified as resistant to *Leishmania*, yet they share other *Leishmania* susceptibility loci within chromosomal regions of T1D susceptibility [81–83].

Subversion of the macrophages as host cells for *L. am.* parasites is essential for the established and persistent infection by the parasites. We have previously demonstrated that during *L. am.* infection in the C57BL/6 mice, OPN is implicated in the strong inhibitory effect of the inflammatory response, exerted by the parasites [35]. In the absence of OPN, this effect is moderated, indicating that OPN is part of the host’s response to the parasites. Infection with *L. am.* of the NOD mice showed an acceleration of T1D onset in the presence of OPN, whilst in its absence T1D decreased and remained lower than in the non-infected animals (Fig. 1B). Therefore, the delay of T1D observed in the absence of OPN after parasite infection indicates that at least in part, OPN-dependent immune mechanisms, known to participate to the autoimmune disease in the NOD mice, are elicited by the parasites [84]. Thus, the protective effect of OPN against T1D is partially abolished by the *Leishmania* infection (Fig. 1B). Moreover, OPN is known to modulate the host response to infection and to be part of immune-mediated inflammation with Th1 cytokine properties in cell-mediated immunity [85, 86]. While Th1 responses are known to be important for parasite clearance in mice, Th2 responses (*i.e*. il-4) are associated with parasite persistence and disease progression [87, 88]. However, human leishmaniasis exhibits mixed Th1/Th2 responses, indicating a more complex immune response in human infections with cutaneous *Leishmania* species compared to mice [89].

Interestingly, in NOD mice *Leishmania* infection highly inhibited *il-17* transcripts (149.5-fold down-regulated at 48h *p.i*., Table 1.1) in BMF, potentially facilitating parasite survival. Indeed, low *il-17* response corresponds to host resistance to *Leishmania* [66]. In contrast, in the absence of opn, *il-17* transcripts increased indicating a role of opn in the suppression of *il-17* in the presence of the parasites and thus participating in the protection against *Leishmania* in the NOD mice. Il-17 has been implicated in T1D regulation in mice, particularly in the later effector phase of the disease [90, 91] and recent reports indicate that il-17 is part of the etiology of T1D [92]. Further research may elucidate the precise mechanisms underlying the interplay between opn, il-17 and T1D in the context of *Leishmania* infection.

Another interesting observation of our data is the variation of the transcripts encoding for il-10. In our experimental setting infection with *L. am.* increases *il-10* transcripts in the BMFs of both NOD wild and *opn* knockout mice, (12-fold and over 50-fold respectively, Table 1.1). Il-10 is an anti-inflammatory cytokine that strongly suppresses Th1-type immune responses [93]. It is involved in the Th1/Th2 immune balance in T1D, while it plays an important role in infections [67, 94]. The absence of il-10 has also been reported to exacerbate both innate and adaptive immunity in response to *Listeria monocytogenes* [95] and plays a role in several other infections [96]. Moreover, il-10 deficiency exacerbates T-cell-mediated autoimmune diseases [67, 94, 97, 98]. In T1D, differential regulation of il-10 was reported and a role in the modulation of both innate and adaptive immune cells and the development of autoimmunity was proposed [94]. This suggests the involvement of OPN in modulating the expression of *il-10* and possibly influencing the immune Th1/Th2 balance. These data corroborate the role of OPN in the efficient development of Th1 immune responses [86].

We assessed the expression of *il-4*, a Th2 cytokine recognized for its role in intracellular pathogens susceptibility when its expression is markedly elevated. It was shown that suppression of il-4 production during early *L. major* infection prevents the differentiation of Th2 responses [99]. It also was reported that il-4 is involved in the initiation of Th1 in the BALB/c mice in response to *L. major*, leading to resistance to the disease [100]. While the role of il-4 on Th2 differentiation is well described, il-4 may also have opposite effects on Th2 immunity [101]. Interestingly, in the NOD BMF infected with *L. am. il-4* transcripts were inhibited independently of the presence of OPN (Table 1.1), suggesting that other pro-inflammatory cytokines known to be potentiated by il-4 may also be absent or inhibited by the parasites in these cells. These data are concordant with the increased proliferation of the parasites in the wild-type BMF (Fig. 5) indicating an OPN-dependent adaptation to the parasites.

OPN in a non-infectious environment is known to repress iNOS expression by increasing ubiquitination and degradation of stat1, the transcription factor for iNOS gene expression [57, 102]. However, the regulation of iNOS expression is complex, contingent upon diverse stimuli, the local cytokine content, and prevailing pathophysiological conditions. One mode of iNOS regulation involves the activation of its promoter by a list of transcription factors, dictating a context-dependent outcome, wherein iNOS expression may be either activated or restrained based on the specific cytokine and microbial stimuli, as well as the cellular context [103, 104]. The data presented revealed a pattern wherein both iNOS and *stat1* are up regulated in the presence of OPN while when OPN is absent, both transcripts decrease (Table 1.1) indicating that OPN plays a role in the upregulation of these genes in this context. Therefore, infection by the parasites activates the iNOS/OPN/STAT1 pathway, suggesting that parasite-induced iNOS activation might contribute to parasite persistence in the macrophages, potentially through the inhibition of the NLRP3 inflammasome by iNOS ([105] and our unpublished observations). Additionally, in the absence of OPN, the low expression of iNOS and *stat1* transcripts, along with the increase of the *il-10* transcripts, indicates a major role of OPN in the Th1/Th2 balance in the NOD mice infected with *L. am.*.

Overall, key findings include: i) differential effects of *L. am.* infection on T1D phenotype between wild-type and knockout mice: while wild-type mice show increased T1D incidence after infection, in the absence of OPN, infection with the parasites decreases the incidence of the disease; ii) higher parasitic load is observed in infected wild-type BMF at 48h *p.i*., while no parasitic proliferation is observed in the NOD *opn* null mice; iii) discrete variations in the *in vivo* clinical phenotype as well as in parasite content at the infection sites are notable in the absence of OPN; iv) pro-inflammatory markers increased after *in vivo* infection in the ear pinna of *opn* null mice at D_100_ *p.i*.. Finally, Th1 responses elicited by the parasites in NOD BMF were abolished in the absence of OPN while Th2 responses were increased.

These findings underscore the intricate interplay between OPN, immune responses, and disease outcomes in the context of both T1D and leishmaniasis, suggesting OPN as a potential target for therapeutic interventions.

## Conclusions

In conclusion, the influence of microorganisms on the development of autoimmune responses poses a complex and intriguing question. Our study investigating the impact of *L. amazonensis* parasites on type 1 diabetes, provides insights into potential immune mechanisms at play in autoimmune-prone NOD mice within an infectious context.

This work serves as an exploratory investigation, and further studies are needed to confirm and expand upon our findings. Additionally, exploring the involvement of genes shared between *Leishmania* and T1D susceptibility loci could offer valuable insights into the interactions between host and parasites in an autoimmune genetic context.

In this context, our findings demonstrate that: a) Environmental triggers, such as infectious microorganisms exemplified by the parasite *L. amazonensis*, which elicits Th1 immune responses like those seen in NOD mice, can impact autoimmunity and b) The interaction between host and parasites may shape singular associations that influence natural selection.

Overall, our study highlights the intricate interplay between environmental factors, immune responses, and genetic predisposition in the context of autoimmune diseases. Further research in this area will contribute to a better understanding of autoimmune pathogenesis and may offer new avenues for therapeutic interventions.

## Materials and Methods

### Mice and ethical statement

NOD/*Shi*LtJ mice were used at 6-8 weeks of age (referred also as wild-type or NOD^+/+^) and were purchased from the Jackson Laboratories (Bar Harbor, ME, United States). *Opn* mutant congenic mice (NOD.B6.Cg.*opn*^-/-^ for simplicity referred to as NOD KO or NOD.*opn*^-/-^) were bred in our facilities [106]. All animals were housed in the animal facility of the Institut Pasteur and all experimental procedures were approved by the Institutional Committees on Animal Welfare at Institut Pasteur (n°: 2013-0014) and carried out under strict accordance with the European guidelines (Directive 2010/63/EU). Animal experimentation was conducted by EM and EG who are both authorized by the Paris Department of Veterinary Services, DDSV.

### Preparation and inoculation of *L. amazonensis* parasites

Parasites were prepared for inoculation following previously described methods [107–109]. Specifically, 10^6^ *L. amazonensis* strain LV79 amastigotes (WHO reference: MPRO/BR/72/M1841) were subcutaneously inoculated into the hind footpad of Swiss nude mice. After 2 months, lesions containing the parasites were excised and purified, according to established procedures [107, 110]. BMF cultures were infected with LV79 amastigotes at a MOI (Multiplicity of infection) of 4:1. LV79 metacyclic promastigotes carrying the firefly luciferase gene in the 18S rRNA locus of *L. amazonensis* LV79 strain nuclear DNA [107] were prepared from amastigotes and cultured at 26°C, as previously described [111]. Infective-stage metacyclic promastigotes were obtained from 6 days old stationary phase cultures by Ficoll gradient and 10^4^ metacyclic promastigotes were injected into the ear dermis of NOD^+/+^ and NOD.*opn*^-/-^ mice. The mice were anesthetized by intraperitoneal administration of ketamine (120 mg/kg Imalgène 1000, Merial, France) and xylazine (4mg/kg; Rompun 2%, Bayer, Leverkusen, Germany). Clinical phenotypes were assessed by evaluating the range of the lesion size on the site of inoculation and compared to the non-inoculated contra-lateral ear as previously described [107]. Ear thickness was measured using a Vernier caliper (Thomas Scientific, Swedesboro, NJ).

### Bone marrow-derived macrophage generation and infections

To obtain macrophages, bone marrow cells were extracted from the tibia and femurs of 6-8-week-old NOD^+/+^and NOD.*opn*^-/-^ mice and differentiated into macrophages (BMFs) using established methods [110]. Briefly, the bone marrow cells were suspended in PBS-Dulbecco enriched with Ca^++^, and Mg^++^, collected by centrifugation and cultured in the presence of rm-CSF-1 (ImmunoTools) at a density of 7.5x10^6^ cells/100 mm Falcon dish and maintained at 37°C (94 % air, 7.5% CO_2_). *L. amazonensis* amastigotes (strain LV79), freshly isolated from footpad lesions of Swiss nude mice as previously described [112], at a MOI 4:1 (parasites: MF) on day 7. The cells were incubated at 34°C for 24 or 48 hours before being lysed for total RNA preparation or placed on glass micro slides fluorescent immunostaining with antibodies against LAMP-1, osteopontin and *Leishmania*.

### Flow cytometry of macrophage-restricted markers

Flow cytometry was used to examine differentiated macrophages from NOD wild-type and knockout mice for the osteopontin gene. Monocyte/macrophage lineage surface molecules including CD11b, CD115, CD11c, MHC-II and the F4/80 antigen were evaluated using the corresponding antibodies [113, 114].

FACS analysis showed that 98% of the cells exhibited similar characteristics, with CD11b^High^, F4/80^High^, and CD115^High^ while surface-specific markers for the dendritic cell (DC) lineage, were negative for CD11c^-^ and high for MHC-II^High^) (Fig. S8). The data was analyzed using Kaluza software on a Gallios flow cytometer from Beckman Coulter [110].

### *In vivo* luciferase-expressing, *L. amazonensis* bioluminescence imaging

Animals were monitored for clinical phenotypes and parasite proliferation at various time points ranging from Day 16 to Day 100. Parasite loads were assessed using bioluminescence imaging as previously described [107]. Briefly, luciferin was injected into animals (*i.p*. 150 mg/kg, D-Luciferin potassium salt, SYNCHEM OHG, Germany) and emitted photons were acquired by a camera from the surface of the entire ear pinna (ROI). The same ROI was examined for all mice at all time points. Total photon emission was expressed in photons/sec/ROI. Median bioluminescence values and SD were calculated for each experimental group. At specific time points, 3 representative mice from each group were sacrificed for further analysis [115]. Contralateral tissues non-injected and control mice groups were analyzed in parallel.

### Immunofluorescence labeling of BMFs

BMFs were cultured on glass slides (CML France) and either infected or not with *L. amazonensis* amastigotes (MOI 4:1) at 34°C (94% air, 7.5% CO_2_). After 24-48 hours, the cells were washed with PBS (Dulbecco), fixed with 4% paraformaldehyde (PFA) for 1h at room temperature and permeabilized with saponin (25 mg/ml). Then, the cells were labeled with primary antibodies as follows: 10μg/ml of the amastigote-specific mAb 2A3-26-biot, and anti-OPN (goat IgG) 7μg/ml (R&D Systems, France) or LAMP-1/CD107a monoclonal antibody specific for the lysosomal-associated membrane protein 1 (LAMP-1) of the parasitophorous vacuole (rat IgG2a FITC, Invitrogen, CA). Revelation was performed with 1.5 μg/ml streptavidin conjugated to Texas Red (Molecular Probes, Cergy Pontoise, France) for the *Leishmania* parasites, with donkey anti-goat FITC (fluorescein isothiocyanate fluorochrome, sc2024, Santa Cruz Biotechnology) for osteopontin and donkey anti-rat-FITC (LS-C351178,

CliniSciences, FR) for LAMP-1. After incubation with the first antibodies (30 min) and three washes with PBS/saponin, a second incubation was carried out for 30 min with the secondary antibodies. Glass slides were then mounted with Hoechst 33342-containing Mowiol 4.88 (Calbiochem), allowing visualization of the DNA of both host cell and amastigote nuclei. Epifluorescence microscopy was used to detect the signals, and the mean protein densities were analyzed by the Zeiss AxioVision Rel. 4.8.2 image acquisition software (Carl Zeiss International). At least 10 different cells were analyzed in at least three different fields. Values are expressed in density units/msec of 20μm^2^ *opn* stained areas/cell. Statistical analyses are performed with the Mann-Whitney test.

### Histopathology

Samples of ear pinna, draining lymph nodes and foot pads were fixed in 4% formalin, embedded in paraffin, and stained with hematoxylin and eosin. Microscopic changes were scored semi-quantitatively by using a five-scale scoring system (1: minimal, 2: mild; 3: moderate, 4: marked, 5; severe) for parameters such as ulceration, acanthosis, necrosis, edema, and presence of parasites. Median values were compared between inoculated and control tissues. Immunostaining with anti-OPN (AF808, R&D systems, Minneapolis, MN, USA) and anti-F4/80 macrophage-specific antibodies (MAB 5580, R&D systems) was performed to estimate the infiltration of neutrophils and macrophages.

### RNA Extraction and Real-Time quantitative PCR

Tissues from representative mice at different time points and 5x10^6^ BMFs were lysed for RNA preparation using the RNeasy Plus Mini Kit (Qiagen, SAS, France) following the manufacturer’s instructions, as described [115]. RNA quality and quantity were evaluated by measuring Optical Density using a Nanodrop ND-1000 micro-spectrophotometer (ThermoFisher Scientific) [116].

Real-time quantitative PCR (qRT-PCR) was performed following the protocol described [115]. The RNAs were reverse transcribed using random hexamers (Roche Diagnostics) and the MMLV-RT reverse transcriptase (Moloney Murine Leukemia Virus, Invitrogen Life Technologies). The relative quantification of the genes of interest was carried out in a 10 μl reaction volume in white ultraAmp 384 well PCR plates (Sorenson, Bioscience, Salt Lake City, UT, USA) using the QuantiTect SYBR Green (Qiagen) and LightCycler® 480 system (Roche Diagnostics, Meylan, France). The primers were used at a final concentration of 0.5 μM (Guaranteed Oligos^TM^, Sigma-Aldrich) and the PCR program consisted of 40 cycles of denaturation at 95°C for 10 sec, annealing at 54°C for 25 sec and extension at 72°C for 30 sec.

The SYBR Green fluorescent emission was measured at the end of the elongation step and crossing Point values (Cp) were determined using the second derivative maximum method of the LightCycler® 480 Basic Software. The qbase program (Biogazelle qbase) for qPCR data management and analysis was used to analyze their Raw Cp values [117]. Eleven candidate control genes were tested as previously described with the geNorm [118] and Normfinder programs [35, 119]. *Hprt* and *ywhaz* were selected as the reference genes for normalizing the gene expression levels (for primers see Table S3). The relative expressions of the interest genes were calculated and *Leishmania* parasites were quantified with the gene target primers (*ssrRNA*) F-CCATGTCGGATTTGGT and R-CGAAACGGTAGCCTAGAG [120].

### Statistical Analysis

The number of parasites and their relative quantification in the BMF cells were determined using the “Mean parasite intensity”, which is the mean number of parasites per infected host cell. The non-infected cells are not taken into consideration. “Prevalence”, on the other hand, is the percentage of infected cells and provides information on the relative sizes of the cells in the study (infected and uninfected).“Crowding” is a measure of the parasites’ density and is defined as the sum of crowding values (parasites living in a cell) divided by the number of parasites [121]. While “mean intensity” is the sum of parasites per infected host, “mean crowding” represents the value of the number of parasites that can live in a host cell on average [122]. To compare parasite load, mean intensity and crowding between macrophages from NOD^+/+^ and NOD. *opn*^-/-^ strains, Quantitative Parasitology (QP3.0) statistical software was used [123]. Mean intensity comparisons were done by a Bootstrap Test [124] and the density-dependent character of parasites was described by Crowding, taking p-values at Cl (Confidence limit 95%) [124].

For the statistical analysis and graphs the GraphPad PRISM T.0 (GraphPad Software, San Diego, CA). A Mann-Whitney test was used to compare transcript relative expression between macrophages from NOD^+/+^ and NOD.*opn*^-/-^ BMFs. The p values were calculated with a Mann-Whitney test to compare ear bioluminescence, ear width, and the transcript relative expression obtained by qRT-PCR between the control group and the group infected with parasites (*: p<0.05; **: p<0.001; ***: p<0.0001).

## Acknowledgements

Our acknowledgments are addressed to the members of the Immunophysiology and parasitism unit who participated in the experimental part of this work.

## Author Contributions

EM conceived and designed the study. EM developed the osteopontin knockout mice in the NOD genetic background and performed the crosses to generate the animals used in this study. EG and EM performed immunofluorescence microscopy of the BMF. LF performed tissue histopathology and interpretation of the histology data. EM cultured cells and performed infections with *Leishmania* parasites *in vitro* and *in vivo*. EG performed and analysed the data of the qRT-PCR. EM wrote the manuscript. All authors participated in the interpretation of the data and read and approved the final form of the manuscript.

## Data availability

The data generated and analysed during this study are included in the core section and the supplementary information of the Additional files. We will make available from the corresponding author upon reasonable request any additional information.

