## Supplemental Tables and Figures for "Type 1 diabetes and parasite infection: an exploratory study in the NOD mouse"

**Table S1A: Representative T1D-related phenotypes**

| | 15-20 wks | 21-25 wks | T1D >26 wks | Mortality >26 wks |
|---|---|---|---|---|
| **NOD Non-INF** | 2/10 (20%) | 4/10 (40%) | 6/10 (60%) | 3/10 (30%) |
| **NOD + *Leish*.** | 2/6 (33%) | 5/6 (83.3%) | 5/6 (83.3% | 4/6 (67%) |
| **KO Non-INF** | 11/20 (55%) | 14/20 (70%) | 16/20 (80%) | 0/20 (0%) |
| **KO + *Leish*.** | 4/12 (33.3%) | 7/12 (58%) | 8/12 (68%) | 5/12 (41%) |

**Table S1B: Impact of *L. amazonensis* infection on T1D in wild-type and *opn* knockout NOD mice**

| | T1D | | INFECTION |
|---|---|---|---|
| | **Non-INF** | **INF** | |
| **NOD$^{+/+}$** | Protection | ++Acceleration | +Proliferation of parasites |
| **NOD.*OPN*$^{-/-}$** | Acceleration | +Acceleration | Non-proliferation |

**Table S2**: **Comparative statistical evaluation of parasite infectivity in BMF**. The statistical analysis as described in the Material section has as follows: "Mean parasite intensity" is the mean number of parasites per infected host cell. The non-infected cells are not taken into consideration. "Prevalence" is the percentage of infected cells and provides information on the relative sizes of the cells in the study (infected and uninfected)."Crowding" is a measure of the parasites' density and is defined as the sum of crowding values (parasites living in a cell) divided by the total number of parasites (Quantitative Parasitology (QP3.0) statistical software). NS: non-significant.

**A. Parasitic infection NOD**

| Host (BMF) | Mean Intensity N° (SD) | | Mean Crowding | | Host size (Infected Cell n°) | |
|---|---|---|---|---|---|---|
| | **24 h *p.i.*** | **48 h *p.i.*** | **24 h *p.i.*** | **48 h *p.i.*** | **24 h *p.i.*** | **48 h *p.i.*** |
| **WT** | 3.64 (2.29) | 12.13 (8.8) | 6.72 | 24.07 | 152 (117) | 58 (54) |
| **KO** | 8.21 (2.4) | 8.5 (2.38) | 9.35 | 8.76 | 29 (0) | 8 (0) |

**B. Parasitic load statistics NOD.** Statistics: Fisher's exact test 2-tailed

| Hosts *p.i.* | Prevalence (P-value) | Mean Intensity (P-value) | Mean Crowding |
|---|---|---|---|
| **WT 24h *vs* 48h** | 0.770 *vs* 0.931 (P=0.0054) | 0.0010 | 6.72 *vs* 24.07 P<0.05, CI 97.5% |
| **KO 24h *vs* 48h** | 1 *vs* 1 (P-value NS) | NS | 9.35 *vs* 8.76 P>0.05 NS |
| **WT 24h *vs* KO 24h** | 0.770 *vs* 1 (P=0.0011) | 0.001 | 6.72 *vs* 9.35 P>0.05 NS |
| **WT 48h *vs* KO 48h** | 0.931 *vs* 1(P-value NS) | NS | 24.07 *vs* 8.76 P<0.05, CI 97.5% |

**TABLE S3**: Sequences of Primers used for qRT-PCR

| **REFERENCE GENES** | **Name** | **FORWARD** | **REVERSE** | **References** |
|---|---|---|---|---|
| *hprt* | *Hypoxanthine-guanine phosphoribosyl transferase* | ATTAGCGATGATGAACCAG | CTTGAGCACACAGAGGG | 1 |
| *ywhaz* | *Tyrosine 3-Monooxygenase/Tryptophan 5-Monooxygenase Activation Protein Zeta* | GTTACTTGGCCGAGGT | GGAGTTCAGGATCTCGT | 2 |
| **TARGET GENES** | **Name** | **FORWARD** | **REVERSE** | |
| spp1 (FL-*opn*) | *Secreted phosphoprotein 1* | TCTGATGAGACCGTCACTGC | CCTCAGTCCATAAGCCAAGC | 1 |
| i-*opn* | *intracellular-osteopontin* | GCCTGTTTGGCATTGCCTCCTC | CACAGCATTCTGTGGCGCAAGG | 1 |
| CD44 | *CD44* | TATCTCCCGGACTGAGG | AGGCATTGAAGCAATATGT | 1 |
| *Il-1$\beta$* | *Interleukin 1 beta* | AGGCAGGCAGTATCAC | CACACCAGCAGGTTATC | 2 |
| *Tbet* | *Th1-specific T-box transcription factor* | AACAAGGGGGCTTCCAAC | TGGCAAAGGGGTTGTTGT | |
| *STAT-1* | *Signal transducer and activator of transcription 1* | TGGGTGCATTATGGGC | TTTCGTGTAGGGCTCC | |
| *IL-4* | *Interleukin 4* | GGAGCCATATCCACGG | AAGCCCTACAGACGAG | 3 |
| *IL-10* | *Interleukin 10* | CCAAGCCTTATCGGAAATG | CCTGAGGGTCTTCAGC | 3 |
| *IL-17* | *Interleukin 17* | CTACCTCAACCGTTCCAC | GCATCTTCTCGACCCT | |
| *IFN$-\gamma$* | *Interferon gamma* | CTTCTTCAGCAACAGCAAGG | TGAGCTCATTGAATGCTTGG | 3 |
| *iNOS-2* | *(Exon 6) nitric oxide synthase 2, inducible* | ACAGGAACCTACCAGC | GGTTGGACCACTGGA | 2 |
| **LEISHMANIA GENE:** *SsrRNA* | *small subunit ribosomal RNA* | CCATGTCGGATTTGGT | CGAAACGGTAGCCTAGAG | 2 |

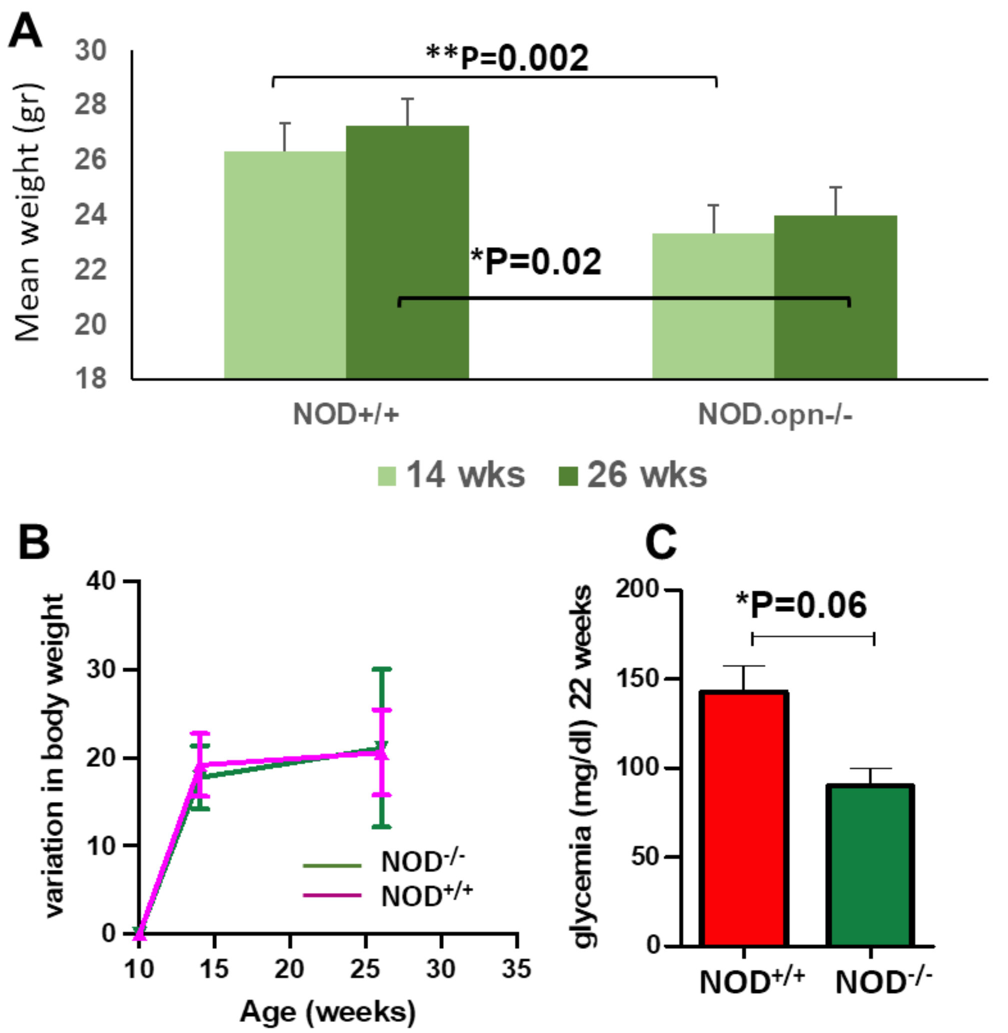


**FIGURE S1**. **Physiology of infection. A.** Mean weight (gr) of the NOD wild-type (NOD$^{+/+}$) and *opn* knockout (NOD.*opn*$^{-/-)}$ mice after *L. amazonensis* infection. **B.** Post-infection variation of body weight, by the age of mice. **C.** Glycemia in infected with *L. amazonensis* NOD wild type and NOD *opn* knockout mice at 22 weeks of age.

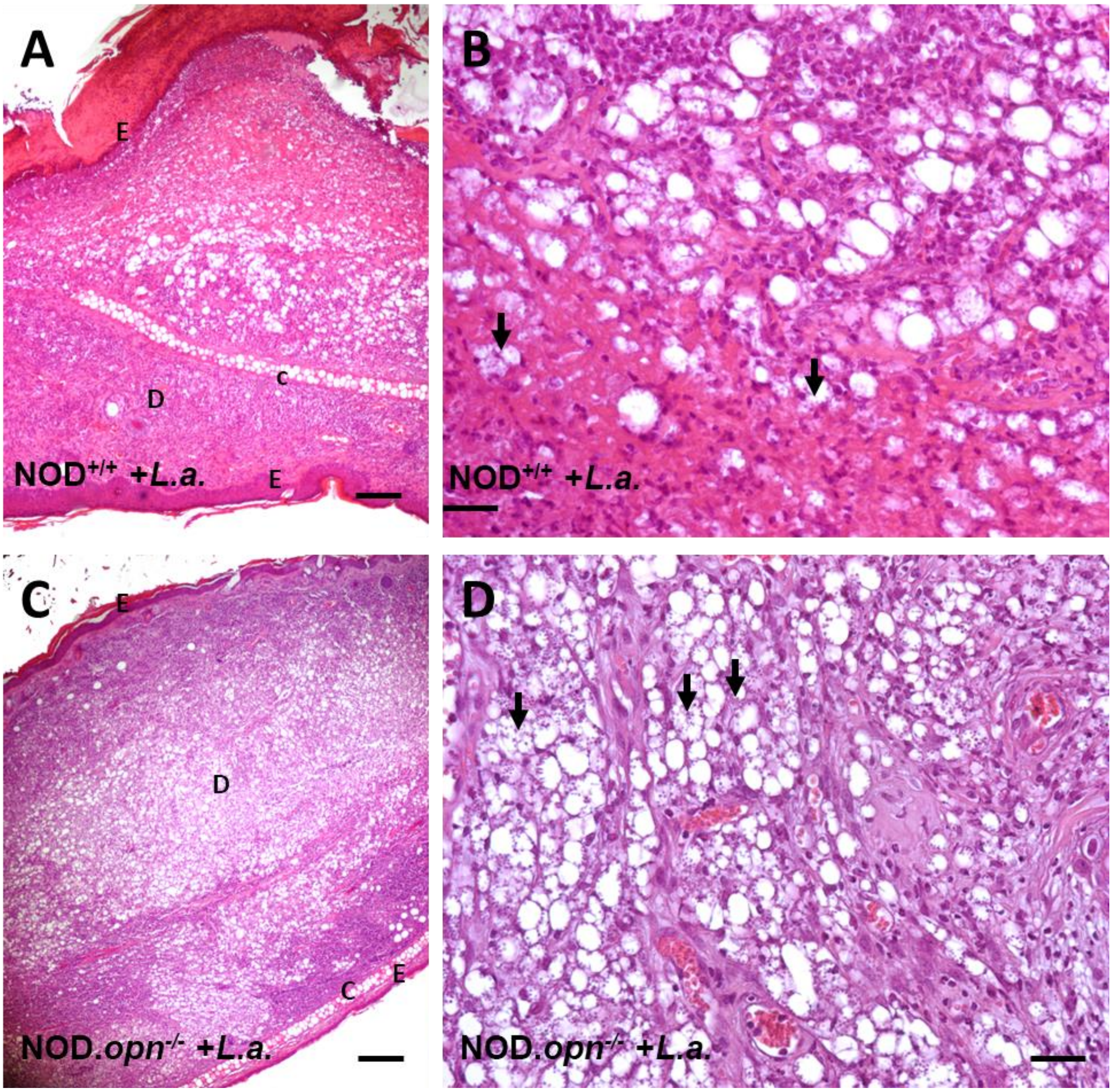


**FIGURE S2: Histological examination of the infected sites in the ear pinna. A & B.** Infected ear pinna from NOD wild-type mice (NOD$^{+/+}$). **C & D.** Infected ear pinna from NOD.*opn*$^{-/-}$ mice. Hematoxylin and Eosin (H&E), **A & C**: Original Magnification x4, scale: 250, **B & D**, Original Magnification x4, scale bar: 50 µm. E: epidermis, D: dermis, C: cartilage. Arrows show the accumulation of parasites in the inflammatory cells.

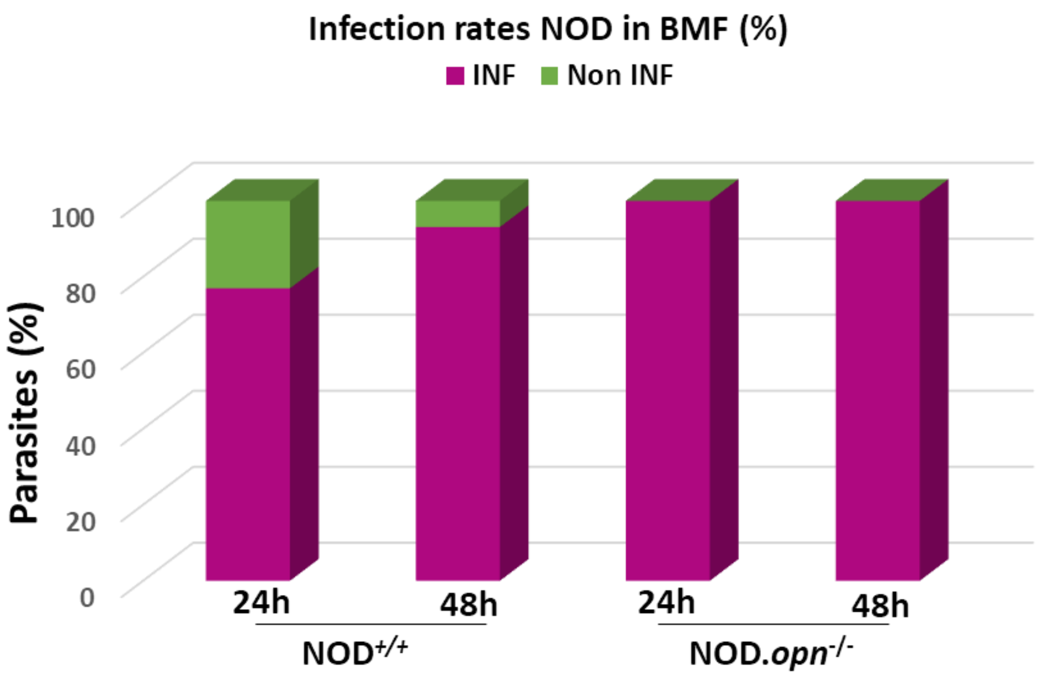


**FIGURE S3. Efficiency of cell transmission of *Leishmania amazonensis* parasites in bone marrow derived macrophages (BMF)**. Representative experiment of BMF infected with with *Leishmania amazonensis,* in the presence (NOD$^{+/+}$) or in the absence (NOD.*opn*$^{-/-}$ ) of osteopontin at 24h and 48h *p.i*. Total cell numbers are presented on Table S2 (QP 3,0 program for stats). NOD$^{+/+}$ at 24h: 152 cells, NOD$^{+/+}$ at 48h: 58 cells. NOD.*opn*$^{-/-}$ at 24h: 29 cells and for NOD$^{-/-}$ at 48h: 8 cells.

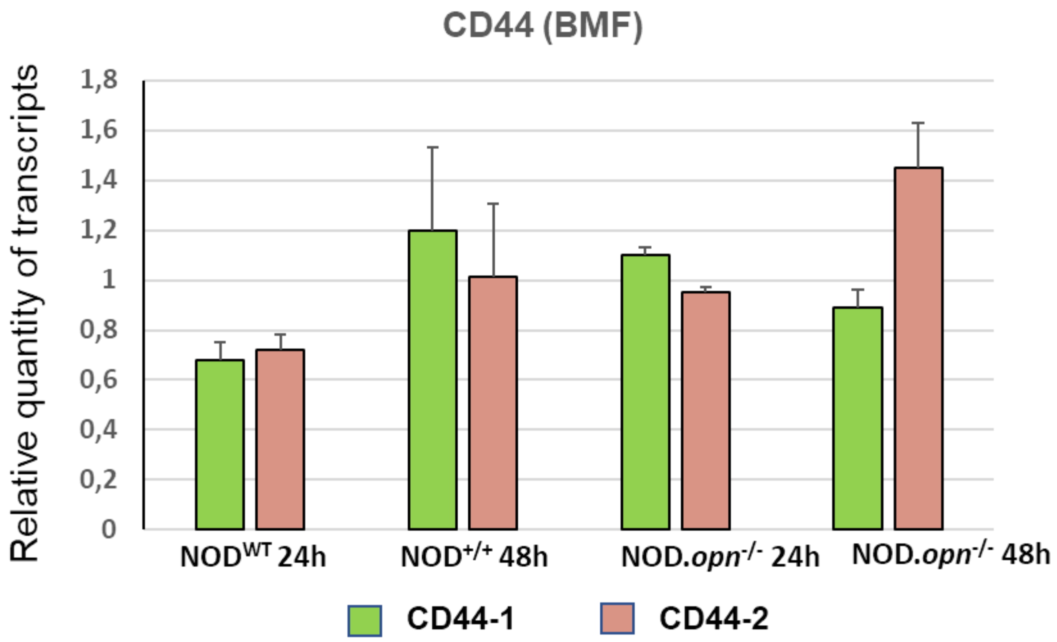


**FIGURE S4. Realtime PCR for the OPN receptor CD44 in the BMF.** Two sets of primers (CD44-1 and CD44-2) were used (see Table S3).

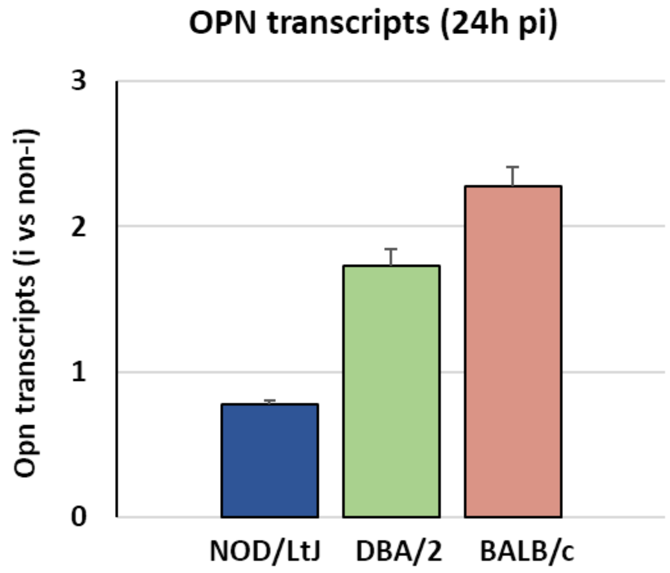


**FIGURE S5**. **Comparison of the relative quantities of *opn* transcripts by q-RT-PCR of NOD/*LtJ*, DBA/2 and Balb/c strains of mice.** Relative quantities are expressed in ratios of infected (i) versus non-infected (non-inf) *opn* transcripts in the BMF at 24h post-infection.

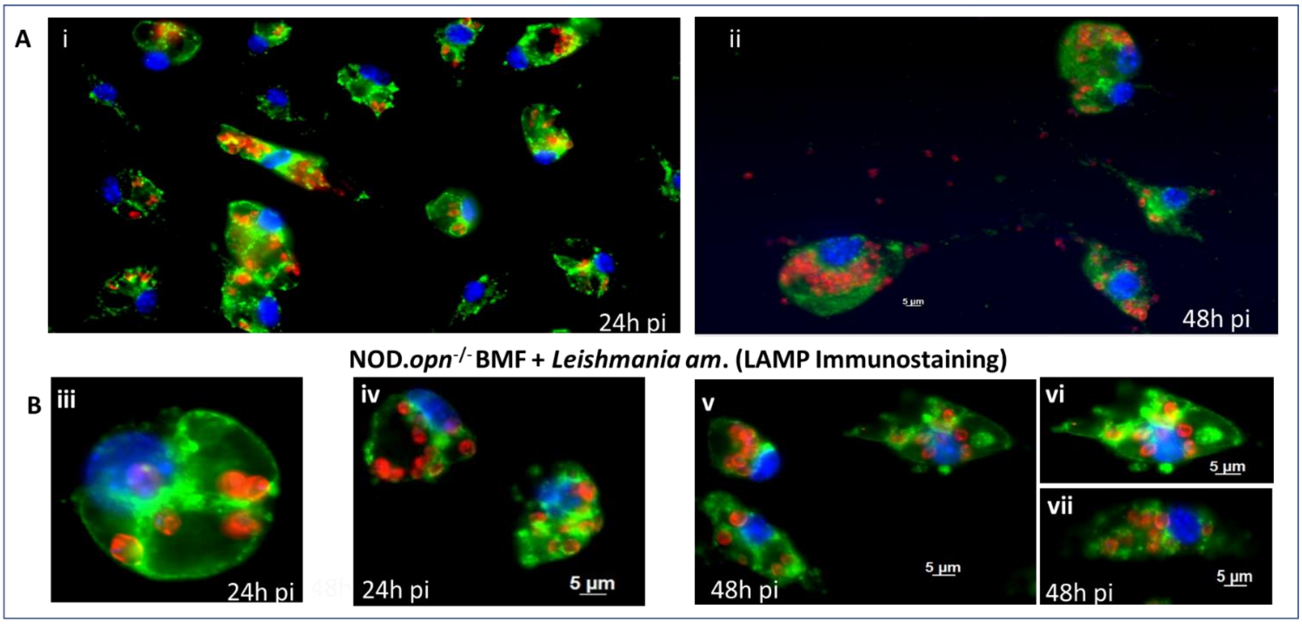


**FIG. S6. OPN favours parasite proliferation in the NOD genetic background (NOD$^{+/+}$). BMF + *Leishmania am*. (LAMP Immunostaining). A.** BMFs isolated from NOD wild-type (NOD/*LtJ or NOD$^{+/+}$*) mice at 24h *p.i.* with *L. am*. (i) and at 48h *p.i.* (ii). **B.** BMFs isolated from NOD KO (NOD.*opn$^{-/-}$* ) mice at 24h *p.i*. (iii and iv) and at 48h *p.i.* (v, vi and vii). Pyroptosis-like cell swelling and membrane blending with intact nuclei and releasing of the parasites were observed only in the presence of OPN (ii). The numbers of cells examined are as in the legend of Fig 7. Parasite Numbers, crowding WT vs KO at 48h: 24.07 vs 8.76; $P<0.05$, CI 97.5%. Data and statistics are from QP 3.0 program as described in Table S2 and in M&M.

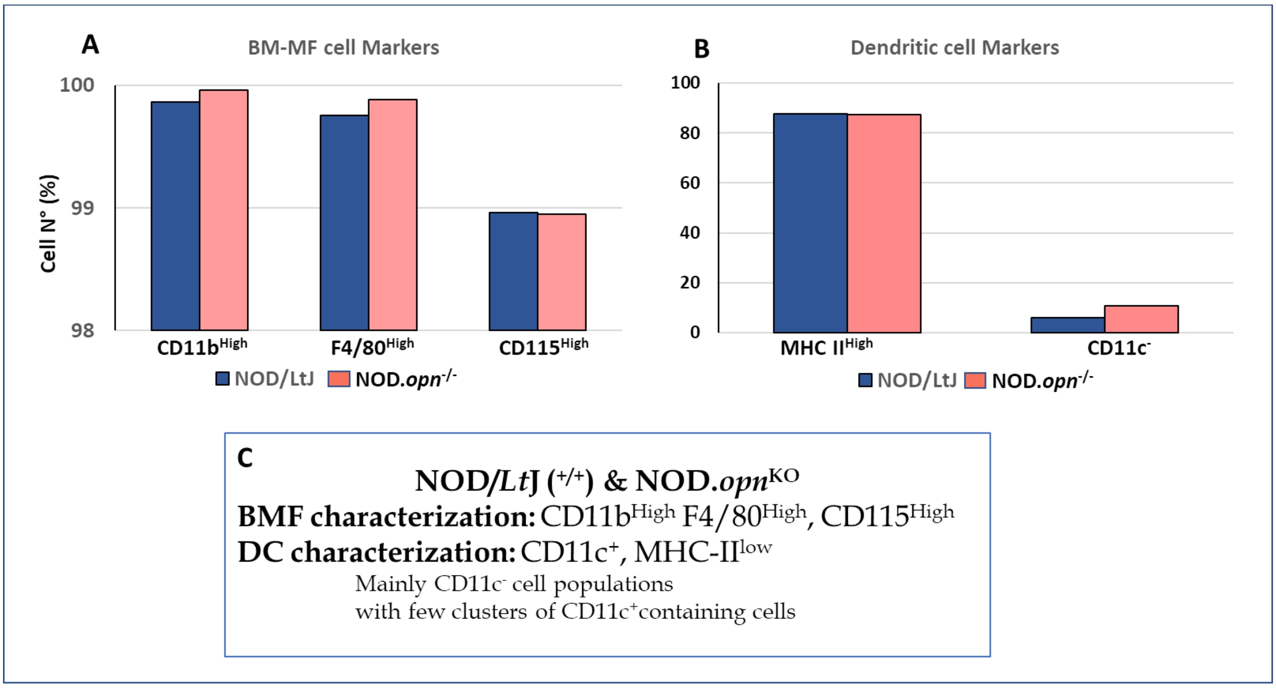


**FIGURE S7**. **FACS analysis of Bone Marrow precursor-derived macrophages (BMF) and Dendritic cells (DC). A.** BMF cell markers and **B.** Dendritic cell markers.

**C.** Summary showing similar characteristics of the BMF and DC between the NOD/*LtJ* (or NOD$^{+/+}$) wild type and *opn* (NOD.*opn*$^{-/-}$) knockout mice. The DC population was MHC-II$^{low}$ and mainly CD11c$^{-}$ with few clusters of CD11c$^{+}$ containing cells.
